# Unravelling human hematopoietic progenitor cell diversity through association with intrinsic regulatory factors

**DOI:** 10.1101/2023.08.30.555623

**Authors:** Patricia Favaro, David R. Glass, Luciene Borges, Reema Baskar, Warren Reynolds, Daniel Ho, Trevor Bruce, Dmitry Tebaykin, Vanessa M. Scanlon, Ilya Shestopalov, Sean C. Bendall

## Abstract

Hematopoietic stem and progenitor cell (HSPC) transplantation is an essential therapy for hematological conditions, but finer definitions of human HSPC subsets with associated function could enable better tuning of grafts and more routine, lower-risk application. To deeply phenotype HSPCs, following a screen of 328 antigens, we quantified 41 surface proteins and functional regulators on millions of CD34+ and CD34- cells, spanning four primary human hematopoietic tissues: bone marrow, mobilized peripheral blood, cord blood, and fetal liver. We propose more granular definitions of HSPC subsets and provide new, detailed differentiation trajectories of erythroid and myeloid lineages. These aspects of our revised human hematopoietic model were validated with corresponding epigenetic analysis and *in vitro* clonal differentiation assays. Overall, we demonstrate the utility of using molecular regulators as surrogates for cellular identity and functional potential, providing a framework for description, prospective isolation, and cross-tissue comparison of HSPCs in humans.

## Introduction

It took more than 50 years from the first experiments for hematopoietic stem and progenitor cell (HSPC) transplantation to become the most successful and widely used form of cell therapy to treat various malignant and nonmalignant hematologic conditions, immune-deficiency illnesses, congenital disorders and to induce tolerance to solid organ allografts ^1,2^. Bone marrow (BM) aspirate was the first source of stem cells, but other types of stem-cell products have been successfully used in the clinic, such as umbilical cord blood (CB) and mobilized peripheral blood (mPB) stem-cell products for HSPC transplantation ^3–6^. When Berenson *et al* demonstrated that BM CD34^+^ cells could reconstitute the hematopoietic system of baboons, CD34 antigen became the hallmark of HSPC ^7^. However, human CD34^+^ HSPCs comprise a heterogeneous mixture of cells representing various lineage commitments and differentiation stages, and are further complicated by age, tissue-specific differences in composition, and phenotype ^8,9^.

Advanced technologies that are able to measure single-cell systems have been used to overcome limitations of bulk cell analysis, providing unique insights into hematopoietic cellular differentiation hierarchy, gene regulatory networks, developmental origin, as well as mechanisms for stem cell aging ^8,10–13^. Single-cell experiments have revealed hematopoiesis is a continuous process rather than a series of discrete states that contain cells of uniform potential, and also have demonstrated that conventional progenitor subpopulations contained previously unappreciated heterogeneity, with varying gene expression states ^14^. Together with pseudotime inference methods, we can also establish a temporal dimension in a static single-cell experiment dataset, capturing the full of the gamut of transitory cell phenotypes ^12,15,16^.

Another important tool to better study the HSPC is the murine model, which from transplantation to transgenics, including humanized mice, has enabled identification of critical pathways in hematopoiesis ^17^. Using time- and tissue-specific genetic deletions, major progress in our understanding of molecular regulation of hematopoiesis was achieved ^17^. Indeed, most of our knowledge of HSPC biology, including important insights into molecular mechanisms that regulates hematopoiesis, has been grounded on murine studies. For example, conditional knockout of *Gata1* in adult mice resulted in the loss of erythroid progenitors and gave rise to a phenotype resembling human red cell aplasia ^18^.

Furthermore, recent scRNAseq data demonstrated an overall strong similarity among the branching structures of hematopoiesis between humans and mice, reinforcing the conservation of molecular networks between mice and man^8^. Despite this, it is widely accepted that human HSPC surface antigen signatures for prospective isolation differ widely from those used for mouse. Given that it is these surface antigens that form the ontology through which we conventionally define the hematopoietic immune system, there exists a shortfall in how we can understand these different cell types and their potential in the human system.

A comprehensive single cell map, merging functional regulators and phenotypic in human HSPCs could guide development and manufacture of better therapies ^19^, ultimately enabling more tailored interventions. Still, to date, most studies human HSPC composition have been compartmentalized, either focusing only on limited surface antigens or HSPC tissue sources at the transcriptome-or epigenome-level ^8,20,21^. We hypothesized that including the single cell quantitation of evolutionarily conserved molecular regulators and transcription factors (TFs) in conjunction with deep single cell phenotypes of human HSPCs would better separate and developmentally order the earliest human hematopoietic cell states. Combined with multiple HSPC tissue sources, this framework could bridge cellular relationships and functional potential(s) across different hematopoietic cell sources to reconcile observed differences and developmental relationships. To accomplish this quantitatively on millions of cells across multiple tissue types and donors, we leveraged single-cell mass cytometry (CyTOF) for highly multiplexed cellular measurement of protein-level targets ^22^. After screening the expression of 328 surface antigens, we optimized the simultaneous capture of 41 protein-level phenotypes and functional regulators on millions of HSPCs, spanning primary human hematopoietic tissues, including adult BM, different conditions of mPB (G-CSF, Plerixafor, or both together), CB, and fetal liver (FL) samples, the major hematopoietic organ during development. We have successfully demonstrated how functional regulators improve the identification and differentiation of cellular identity and functional potential. We present several key implications: 1) our results strongly support the direct origin of both erythroid and megakaryocyte lineages from HSCs; 2) we advanced the understanding of myelopoiesis branching and provide clarity regarding lineage commitments; 3) our investigation revealed a higher frequency of early HSPCs in both mPB and CB when compared to BM. Additionally, we identified a diverse range of unique cellular phenotypes specifically associated with fetal liver (FL). Overall, we reveal new insights into the earliest events of human hematopoietic immune development and provide new definitions for prospective isolation of HSPCs and comparison of HSC and HSPCs across different tissues in humans.

## Results

### Single cell molecular regulator expression refines the phenotypic landscape of human BM progenitors

To build up a deep phenotypic antibody panel for human HSPCs, we firstly performed a flow cytometry screen with 328 surface antibodies on CD34-enriched healthy human BM (n=2), enabling discovery of new surface markers specific for HSPCs based on differential expression across progenitor populations (Fig. S1A-B). Based on the results of this screen, we observed that CD84, CD164, and CD172ab differentially tracked across maturation and were therefore incorporated into our subsequent cytometry panel. We stained healthy adult CD34-enriched BM mononuclear cells (BMMNC, n=8) with a CyTOF antibody panel targeting HSCs, progenitor, and hematopoietic lineages, together with known TFs and molecular regulators essential and conserved in human and murine hematopoiesis, totaling 41 protein-level targets (Table S1). This dataset consisted of approximately 1 million CD34+ and 2 million CD34-cells (Fig. 1A).

**Figure 1:**
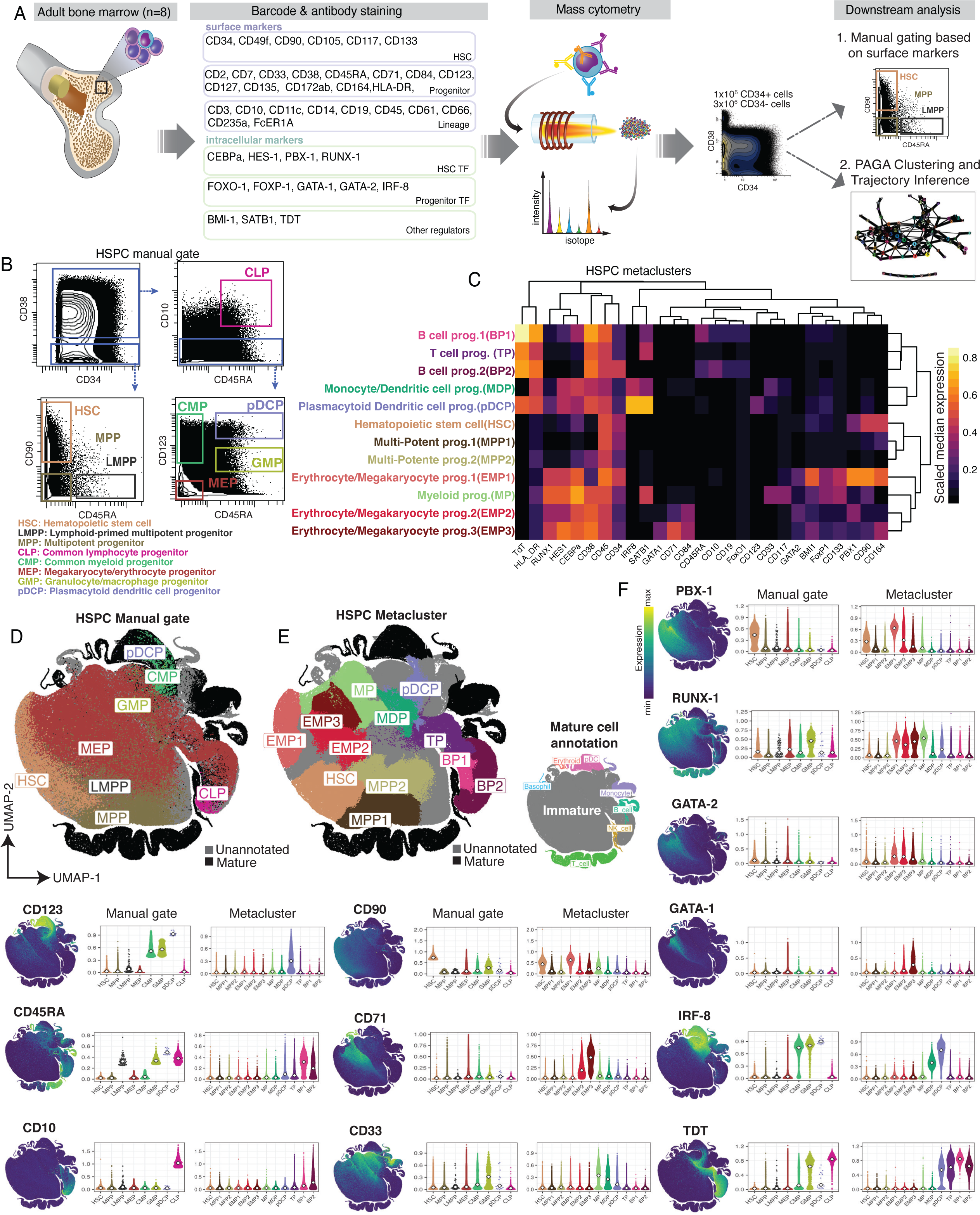
The combination of surface markers and molecular regulators refines the phenotypic landscape of human BM progenitors. A) Human adult bone marrow mononuclear cells (BMMNC) were barcoded and antibody stained for mass cytometry and the resulting data were 1) manually gated and 2) analyzed using the PAGA pipeline for clustering and trajectory inference. B) Gating strategy for conventional HSPC populations. C) Heatmap of median expression of HSPC metaclusters. D) UMAP of CD45+ BMMNCs colored by manual gate. E) UMAP colored by HSPC metaclusters (left) or mature metaclusters (right). F) UMAPs colored by protein expression (left) alongside violin plots of the same protein separated by manual gate (center) and metacluster (right). Diamond indicates median.

After pre-processing and batch correction (*Methods*; Fig. S2A-B), we manually gated cells based on canonical HSPC schemes using phenotypic surface markers ^23^ (Fig. 1B). While updates to this standard model of hematopoiesis have been suggested ^9^, we used it here as an operational paradigm for to maximize comparability across studies. In parallel, we performed leiden clustering ^24^ and partition-based graph analysis (PAGA) ^25^ on CD34+ and CD34-cells using all protein targets (*Methods*). Conversely, the clusters here were annotated with cell population labels based on their molecular phenotypic identity (Table S2, Fig. 1C, S2C). For instance, the expression of CD33, CD123, and HLA-DR together with the molecular regulators, such as TdT and IRF-8 were crucial for the annotations of the Monocyte/Dendritic cell progenitor (MDP) cluster and the plasmacytoid dendritic cell progenitor (pDCP) cluster. TdT was also useful for annotation of B and T cells progenitors (BP1, BP2, and TP), as previously described ^8,26^. We annotated two multipotent progenitor populations (MPP1 and MPP2) in the early HSCP compartment (CD34+CD38-/lo cells), separated primarily by a slight upregulation of CD38 in MPP2. Although MPP is the immediate progeny of HSC, we observed downregulation of molecular regulators, such as PBX-1, CEBPa and BMI-1, in the differentiation of HSC to MPP ^27^.

By projecting cells onto a Uniform Manifold Approximation and Projection (UMAP) plot ^28^, we observed a substantial improvement in population separability in the metaclustered HSPC subsets, as compared to manually gated populations (Fig. 1D-E). For example, in the manually gated data, megakaryocyte/erythrocyte progenitors (MEPs) are diffusely spread across the UMAP, reflecting overlapping phenotypic expression profiles with early, myeloid, and lymphoid progenitors. In contrast, the erythrocyte/megakaryocyte progenitors (EMPs) in the metacluster projection localize in a concise region on the UMAP, reflecting their more unique phenotypic and molecular expression patterns.

The expression of several important surface and molecular regulators enabled discernment of HSPC populations (Fig. 1F, S2D). We observed high expression of the transferrin receptor, CD71, restricted to EMPs aligning with recent literature reporting CD71 expression in erythroid progenitor ^9,29^. We also observed higher expression of IRF8 restricted to myeloid and pDCP metaclusters, in agreement with previous studies reporting that IRF8 was essential for the development of monocytes and dendritic cells ^30,31^. Taken together, combining surface phenotype with expression of key molecular regulators resulted in a more coherent definition of BM HSPCs and detection of novel putative HSPC identities.

### Trajectory analysis reveals a direct HSC to erythrocyte/megakaryocyte lineage commitment arm

Based on our clustering analysis, we identified three putative populations with erythroid/megakaryocyte commitment: EMP1 that belongs to the early HSPC compartment, due to lower expression of CD38, and EMP2 and EMP3, both of which are in the CD34+/CD38+ progenitor compartment (Fig. 1C). TFs such as RUNX1, FOXP1, PBX1, GATA1 and GATA2, together with the expression of CD71 were important to guide these annotations. Interestingly, we observed that expression of CD84, a member of CD2 subfamily of the immunoglobulin receptor superfamily ^32^, was restricted to EMP2 and EMP3, while CD164, a sialomucin, was expressed only in the EMP1 and EMP3 populations (Fig. 1C).

To better define the differentiation trajectory of erythrocytes, we limited our analysis to only include cells from connected BM clusters representing HSC, EMP1-3, and CD34-CD235ab+ immature erythrocyte populations (*Methods*). We then reclustered and regraphed these data using only surface markers and molecular regulators expressed by these populations. This approach yielded a direct and unbranched trajectory from HSC to EMP1, EMP2, EMP3, and then immature erythrocytes (Fig. 2A). Interestingly, addition of MPPs to this analysis resulted in a spurious branch containing only MPPs, suggesting MPPs are not part of the erythroid trajectory (Fig. S3A).

**Figure 2:**
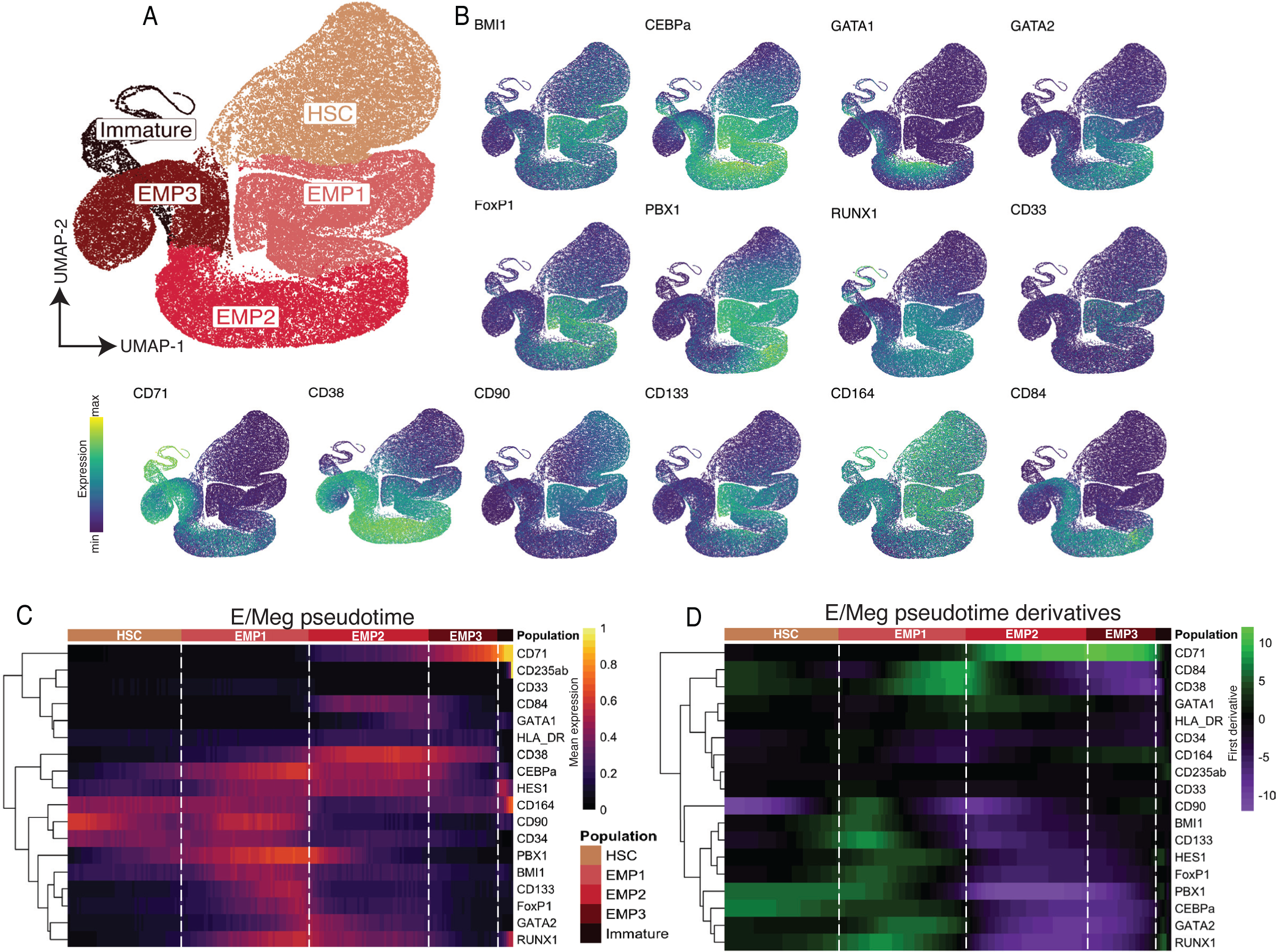
Trajectory analysis reveals a direct HSC to erythrocyte/megakaryocyte lineage commitment arm. A) UMAP of BM cells in the erythroid trajectory, colored by metacluster. B) UMAP colored by marker expression. C) Heatmap of mean expression along the erythroid/megakaryocyte pseudotime trajectory. D) Heatmap of first derivative of marker expression, showing rate of expression change, along the pseudotime trajectory.

As we observed in the larger HSPC compartment, the combination of surface marker and molecular regulator co-expression patterns defined and separated progenitor populations (Fig. 2B). EMP1s were characterized by co-expression of RUNX1, GATA2, and PBX1, while EMP2s upregulated expression of GATA1, CD38, CD84, and CD71. EMP3s downregulated molecular regulators and CD38, while increasing expression of CD71. Organizing cells into a pseudotime axis in an unsupervised manner facilitated the ordering of expression patterns and corroboration of the putative differentiation trajectory from HSC to immature erythrocyte (Fig 2C). We observed PBX1 expression in the early progenitor compartment prior to expression of GATA2, which was then followed by expression of GATA1. Additionally, we observed some co-expression of GATA1 with RUNX1, which may be indicative of megakaryocytic potential in those cells^33^.

To quantify the rates of change in expression values and to identify inflection points between up- and downregulation of proteins, we calculated the derivative of protein expression across the pseudotime axis (Fig. 2D). While we found substantial acceleration of molecular regulator expression in EMP1s, upregulation of several of these proteins, including RUNX1 and PBX1, began in the HSC compartment, suggesting that some HSCs were already biased towards an erythrocyte/megakaryocyte fate. Interestingly, we observed a stark inflection point at the border between EMP1 and EMP2. This inflection point is characterized by coordinated downregulation of RUNX1, GATA2, and PBX1, with simultaneous upregulation of CD71. In summary, we observed direct emergence of erythroid/megakaryocyte progenitors from HSCs, characterized by expression of several lineage-specific molecular regulator and surface markers.

### Epigenetic characterization of EMPs confirms a continuous increase in erythroid and megakaryocyte molecular lineage commitment

Epigenetic priming of transcriptional programs represents a hallmark of lineage commitment in developmental and stem cell biology. To assess relevant TF signatures in our putative EMPs, we prospectively isolated EMP populations by fluorescence-activated cell sorting (FACS), applied Fast-ATAC to measure genome-wide chromatin accessibility, and modeled TF binding dynamics through accessibility at motif sites with the TOBIAS algorithm ^34–37^ (Fig. 3A and S3B-C). We obtained high quality ATAC seq data with a mean ± standard deviation TSS score of 14.5±1.08, across our biological and technical replicates (Fig. S3D-F). When comparing between our sorted populations, the most significantly enriched TF motifs in EMP1 and EMP2 populations were involved in HSC developmental and maintenance, such as EGR1, KLF15, and members of the TCF family (Fig. 3B) ^38–40^. Additionally, the early erythroid-megakaryocytic differentiation factor TCF4 motif was also highly enriched earlier in erythroid differentiation ^41^. EMP3, on the other hand, presented the most erythroid/megakaryocyte committed epigenetic profile, with the highest accessibility of the GATA TF family and MECOM ^42,43^. Furthermore, EMP3 exhibits partial overlap with a high-GATA-1-expressing BM progenitor cells we previously identified, sharing similarities in terms of both epigenetics and functionality with erythroid progenitors involved in human red blood cell development ^44^.

**Figure 3:**
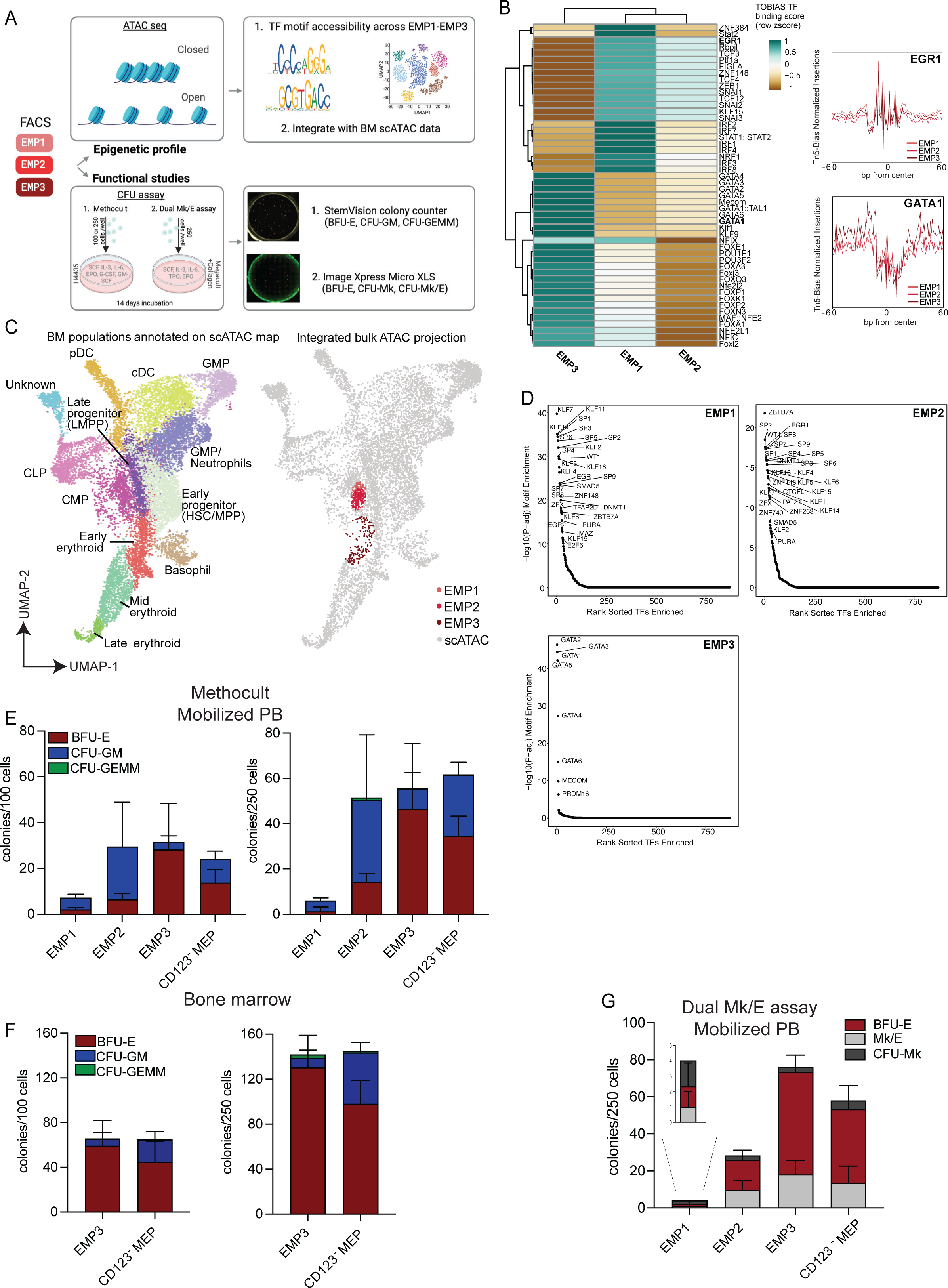
ATAC-seq and *in vitro* clonal differentiation assays of erythrocyte/megakaryocyte progenitors corroborates lineage commitment. A) Experimental pipeline: EMP populations were sorted by FACS and subject to ATAC-seq and epigenetic analysis (top) and *in* vitro functional CFU assays (bottom). B) Heatmap of the binding z-scores of the top 50 transcription factors significantly enriched within EMP states (left), and bias-corrected, normalized Tn5 insertions across all EGR1 and GATA1 motif sites in consensus peaks (right). C) UMAP previously published and annotated BM scATAC dataset with Seurat clusters manually annotated colored by cell population (left) and projection of the EMP populations ATAC-seq data simulated as scATAC counts onto scATAC UMAP space (right). D) Dotplot of TF enrichment p-value ordered by rank sorted enrichment for each EMP population. E) Methylcellulose assay. Barplots quantifying the number of colonies formed per 100 input cells (left) or 250 input cells (right) of sorted mPB erythroid populations, colored by colony lineage. Error bars represent standard deviation (s.d.) F) Same as E) for BM. G) Same as E) for dual Mk/E assay. Abbreviations: BFU-E: burst forming unit-erythroid; CFU-GM: colony forming unit-granulocyte and monocyte; CFU-GEMM: colony forming unit-granulocyte, erythroid, monocyte, and megakaryocyte.

To assess how the epigenetic profiles of our EMP populations related to the HSPC landscape, we projected our data onto a publicly available scATAC dataset of BMMNCs^45^. We observed that EMP1 and EMP2 were epigenetically similar to one another and projected at the interface of the previously annotated CMP and early Erythroid populations, whereas EMP3 projected onto the erythroid arm of the scATAC UMAP (Fig. 3C, S3G). To better understand TF dynamics in our sorted populations within the context of whole BM, we looked at TF motif enrichment of our single-cell simulated EMP1-3 populations compared to all other annotated BM populations (Fig 3D). We observed high accessibility at TF motifs of KLF, EGR and SP families for EMP1 and EMP2 populations, while EMP3 presented an exclusive enrichment for E/Meg related TF motifs (Fig. 3D). Taken together EMP1 and EMP2 populations still presents non-erythroid and non-megakaryocyte features in their epigenetic profiles, retaining features of multi-potent HSPC progenitors, while EMP3 seems to be more restricted to E/Meg lineage. Furthermore, these epigenetic data further support our single cell proteomic trajectory (Fig. 2A-D) of EMPs emerging from the HSC compartment and differentiation directly towards an erythroid and megakaryocyte restricted state.

### Functional, clonal characterization of erythrocyte/megakaryocyte progenitors corroborates lineage commitment

To assess if the lineage potential of our EMPs matched their proteomic and epigenetic profile, we FACS purified EMP1, EMP2, EMP3, as well as the canonical CD123-MEP cells from two mPB samples and performed a colony-forming unit (CFU) assay in MethoCult (Fig. 3A and S3B-C). We found that all four populations differentiated into erythroid burst forming units (BFU-E), with a gradual increase in clonal output from EMP1 to EMP3 (Fig. 3E). EMP3 had the purest capacity for erythrocyte differentiation, even compared to CD123 negative MEPs. EMP1 and EMP2, however, also yielded many CFU-GM colonies, confirming the presence of non-erythroid and non-megakaryocyte program. While the EMP3 population was also not 100% pure for erythroid differentiation, the frequency of BFU-E in our data (>80%) is in alignment with a recent report on human megakaryocyte-erythroid progenitors ^46^. The consistency of these results was also confirmed in adult BM (Fig. 3F).

Sanada *et al.* described a novel dual-detection functional *in vitro* CFU assay for single cells that enables differentiation into both megakaryocyte and erythroid lineages^46^. We applied this approach to our four sorted populations from two mPB donors and observed a gradual increase in the total number of colonies from EMP1 to EMP3, similar to our observations in Methocult. EMP1, EMP2, and EMP3 presented a mixed potential for all three colony types, BFU-E, CFU-Mk, and CFU-Mk/E, with EMP3 presenting an enrichment for BFU-E (Fig.3G). These functional data validate our hypothesis that the combination of surface with intrinsic molecular regulators results in enhanced separation, identification, and subsequent purity of specialized HSPC subsets. Moreover, our data supports the idea of erythrocyte and megakaryocyte lineages emerging from multipotent progenitor cells directly^9^ and provides a high-resolution single cell map to further interrogate in the future.

### Trajectory analysis refines myelopoiesis branching and lineage commitments in HSPCs

To better define the differentiation trajectory of mononuclear, myeloid cells, we employed the same computational pipeline used for the E/Meg lineage. We limited our analysis to only include cells from connected BM clusters representing early progenitors, myeloid progenitors, and mature myeloid cells (*Methods*). We then re-clustered and re-graphed these data using only surface markers and molecular regulators expressed by these populations. This refined myelopoiesis graph revealed a differentiation pathway originating from HSC, continuing through MPP, MP, to MDP (Fig. 4A). Here, the trajectory branched, with one arm continuing to pDCP and pDC, and the other differentiating to a monocyte/conventional dendritic cell progenitor (McDP), before branching again to mature monocytes and cDCs. The MDP, which served as a junction for all three lineages, was defined by co-expression of CEBPa, RUNX1, and IRF8 (Fig. 4B). Arranging each lineage into its own pseudotime trajectory revealed dynamic of expression of CEBPa, a key regulator of myeloid and dendritic cells ^47,48^ (Fig. 4C-E). Derivative analysis of the pseudotime trajectories revealed a high rate of IRF-8 upregulation at the MDP branch point, across all three lineages (Fig. S4A-C). We also observed transitory expression of TdT in the pDC lineage, which may reflect rearrangement of the immunoglobulin heavy chain locus ^49^.

**Figure 4:**
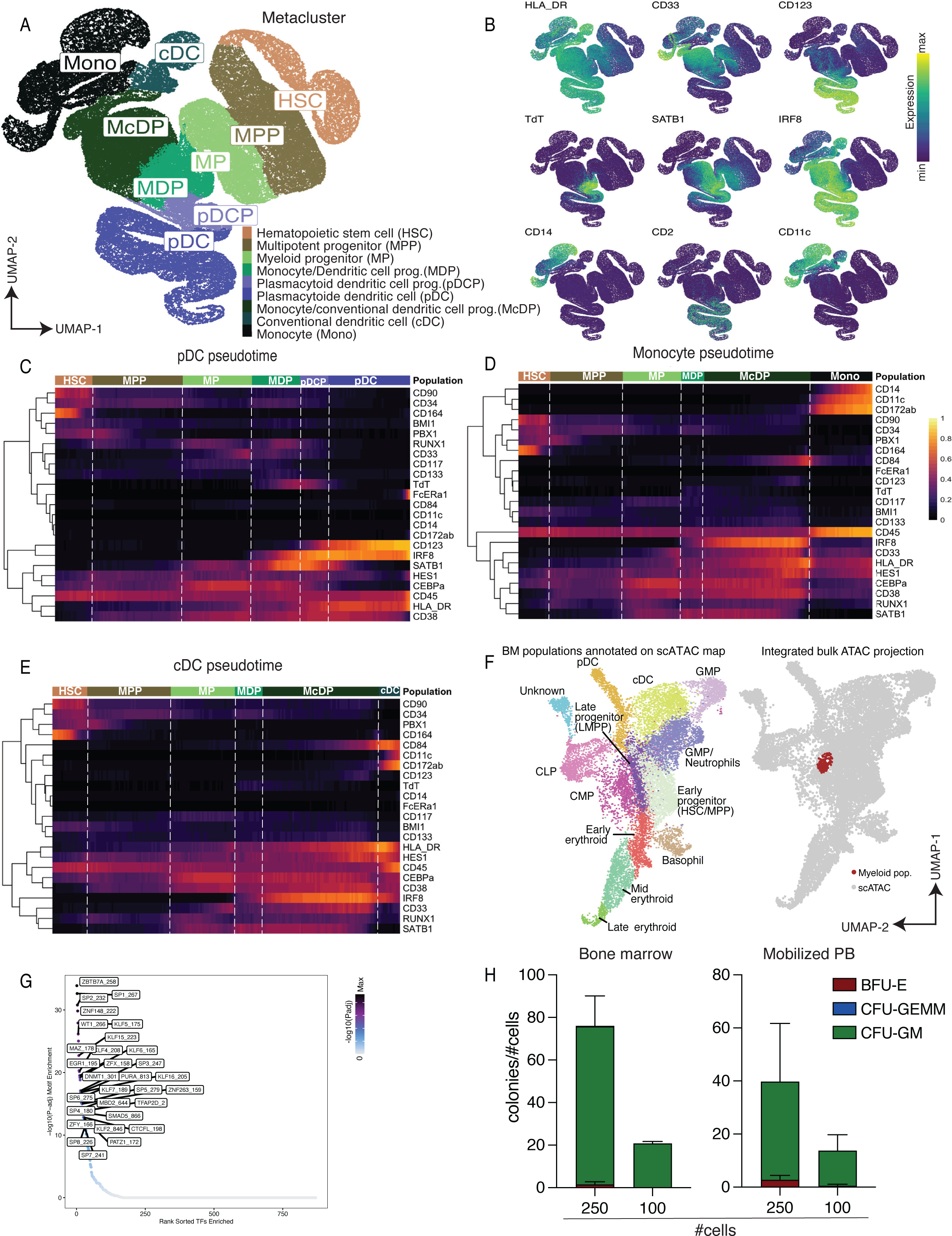
Epigenetic and functional characterization of myeloid progenitors corroborates lineage commitment. A) UMAP of BM cells in myeloid trajectories, colored by metacluster. B) UMAP colored by marker expression. C) Heatmap of mean expression along the pDC pseudotime trajectory. D) Same as C) for the monocyte trajectory. E) Same as C) for the cDC trajectory. F) UMAP previously published and annotated BM scATAC dataset with Seurat clusters manually annotated colored by cell population (left) and projection of the sorted myeloid population ATAC-seq data simulated as scATAC counts onto scATAC UMAP space (right). G) Dotplot of TF enrichment p-value ordered by rank sorted enrichment for sorted myeloid ATAC data. H) Methylcellulose assay. Barplots quantifying the number of colonies formed per input cell number of sorted myeloid population, colored by colony lineage from BM (left) and mPB (right). Error bars represent s.d.

Unlike the case of the EMPs, these myeloid progenitor population were best defined by intracellular molecular regulators, therefore it was not possible to prospectively isolate all populations for functional assessment of lineage potential (Fig. 4 A-E). Still, we were able to come up with a scheme to isolate an overlap of the MP, MDP, and McDP myeloid progenitor subsets *in vitro*, based on CD34+CD38+CD71-/loCD133+CD84-CD33+(Fig. S4D-E). To assess their epigenetic landscape, we again performed Fast ATAC-seq on two technical replicates of these cells from mPB (Fig. S4F-H). When we projected the Fast ATAC-seq data of our myeloid population onto the scATAC UMAP of BMMNCs ^45^, we observed co-localization within the conventional CMP and late progenitor populations (Fig. 4F, S4I). Further TF motif enrichment across BMMNCs showed increased activity of myeloid and monocyte TFs such as SP and KLF factors (Fig. 4G).

In parallel, we performed CFU in Methocult of the sorted myeloid population to assess the differentiation potential of these cells. We found that regardless of HSPC source, BM or mPB, the sorted myeloid population yielded highly pure CFU-GM colonies, confirming the strong myeloid signature of these cells (Fig. 4H). In summary, we refined myelopoiesis into more granular populations using a combination of surface markers and molecular regulators and experimentally confirmed their myeloid epigenetic profile and lineage commitment.

### mPB samples are highly enriched for early HSPCs

Despite the research focus on bone marrow aspirates and cord blood as human study materials, mobilized peripheral blood now represents the predominant source for therapeutic application of HSPCs in the adult (source Center for International Blood and Marrow Transplantation Research – CIBMTR). One of the first mechanisms of action described for G-CSF mobilization of HSPC was the impairment of the SDF-1/CXCR4 interactions ^50^. Later, Plerixafor (AMD3100), an antagonist of the chemokine receptor CXCR4, that was initially developed as a treatment for HIV, was also shown to induce rapid mobilization of HSPC, offering an alternative to G-CSF mobilization ^51^. It has been almost 30 years since HSPC mobilization was first used in a clinical setting, yet there are relatively limited studies on the graft HSPC composition or the effect of different mobilizing agents ^52,53^. To extend our deep phenotypic analysis of HSPCs beyond BM, we applied our mass cytometry experimental and analytic pipeline to several mPB samples induced under different mobilized conditions: five G-CSF mPB samples, three Plerixafor (Plex) mPB samples, and one G-CSF combined with Plex (G+P) mPB sample.

To characterize and compare HSPCs in BM and mPB, we gated CD34+ cells from each tissue and performed Leiden clustering. To compare the overall difference in cell composition between samples in an unbiased manner, we quantified the pairwise Manhattan distance between all samples, based on Leiden cluster proportions, and hierarchically clustered the resulting distance matrix. While the majority of BM samples clustered together, several BM samples clustered with the mPB samples and the mPB samples did not cluster by mobilization condition, underscoring the heterogeneity in cell composition of both BM and mPB samples (Fig. 5A).

**Figure 5:**
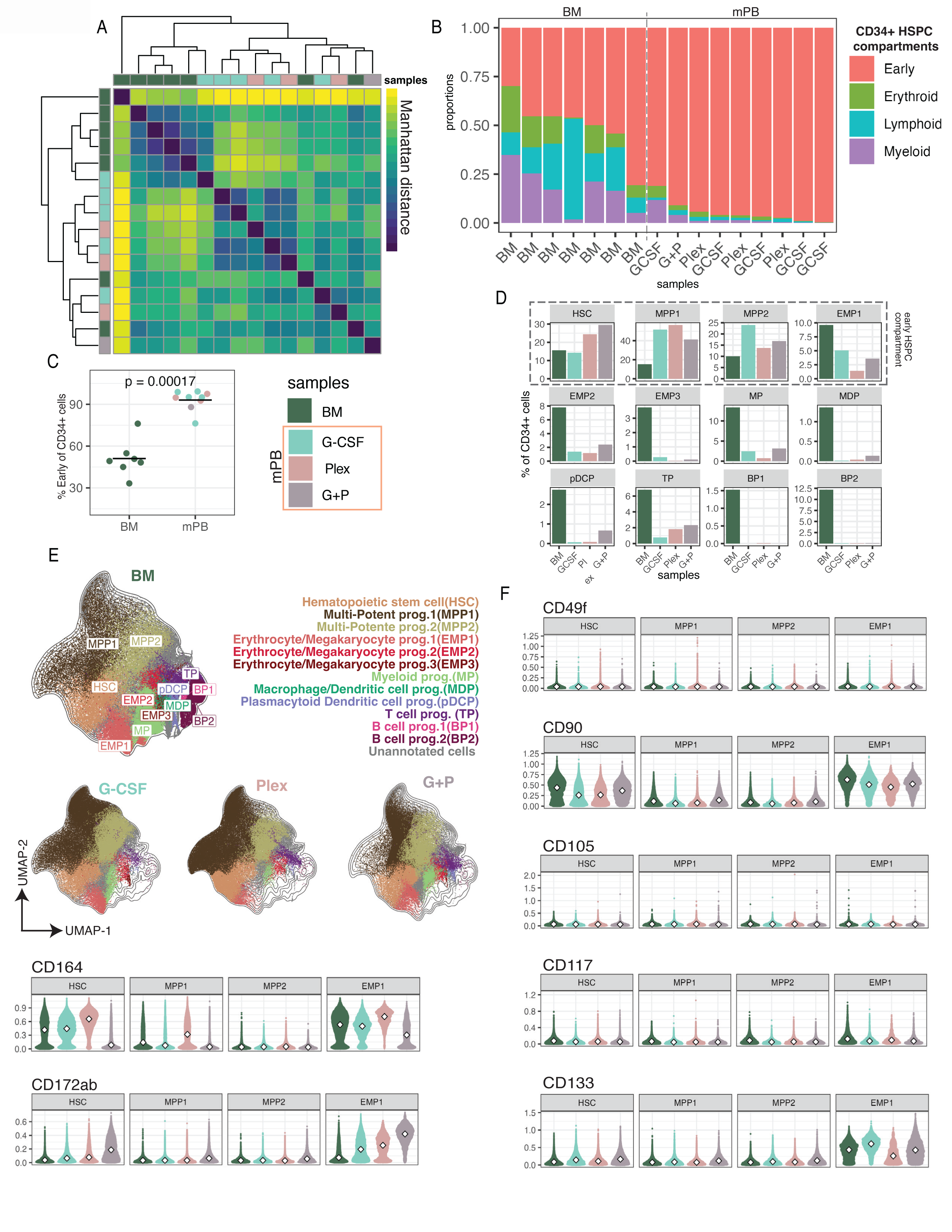
mPB samples are highly enriched for HSPCs. A) Heatmap of Manhattan distance between donor samples based on leiden cluster frequencies of CD34+ cells, colored by tissue/product (left and top annotation colors). B) Barplots quantifying frequency of cells in CD34+ HSPC compartments based on leiden clustering (colors) by donor samples (bars), labeled by tissue/product. C) Dotplot quantifying frequency of Early HSPCs progenitors by tissue/product. P-value derived from Wilcoxon rank sum test. D) Barplots quantifying frequency of HSPC metaclusters by tissue/product. E) UMAP of BM and mPB CD34+ cells colored by metacluster and separated by tissue/product. F) Violin plots of marker expression by tissue/product. Diamond indicates median.

BM cells labeled with their original metacluster annotations (Fig. 1) were used to train a classifier to automatically assign metacluster annotations to phenotypically similar mPB cells (see *Methods*). To assess the overall HSCP composition of each tissue, we grouped annotated cells into four primary lineages, “early”, “lymphoid”, “myeloid”, and “erythroid”, and quantified their frequencies for each sample (Fig. 5B). Arranging samples by increasing levels of early HSPCs perfectly separated BM from mPB samples (Fig. 5B), with significantly greater representation in mPB (Fig. 5C; p=0.00017). We confirmed this significant enrichment for early HSPCs in mPB samples by manually gating CD34+CD38- and comparing tissues (Fig. S5A; p=0.00035), supporting findings in the literature in which G-CSF and Plex mobilize more early HSPC ^52^. This enrichment was most pronounced in HSC, MPP1, and MPP2 populations in all three mobilization conditions (Fig. 5D). UMAP visualization of cells from all BM and mPB samples illustrated that the expansion of the early HSPC compartment in mPB came at the expense the more committed HSPC clusters when compared to BM cells, particularly for clusters implicated in lymphoid progenitor commitment (Fig. 5E).

Since our classifier for assigning metacluster IDs has some flexibility from the original BM-based definition, we then asked if there were tissue-or mobilization-specific differences in expression within our metacluster annotations. We visualized expression of several key markers for the identification of human HSC, such as CD133, CD105, CD164, CD49f, CD117, CD90 across samples within the early HSPC compartment (Fig. 5F) ^8,54–57^. The most drastic differences were observed in the expression of the sialomucin protein CD164. Possibly playing a role in adhesion, CD164 has been suggested as an alternative to overcome the shortcomings of CD38-based gating strategies in HSPC transplantation settings ^8^. Although we analyzed only one sample for the G+P mobilization condition, we observed an increased expression of CD172ab, a member of the Signal Regulatory Protein (SIRPα) family, in the HSC population of this sample, as well as in the EMP1 and MP metaclusters (Fig. S5B). CD172ab expression was previously observed on immature CD34+/CD38-hematopoietic cells, but it is also a myeloid-specific immune checkpoint ^58^. Finally, we observed differences in the expression of some surface markers and molecular regulators across the subpopulations in the CD34+/CD38+ compartment, such as CD33, CD45RA, CD123, GATA1, FOXP1, although the functional implications of these differences are unclear (Fig. S5B-C).

Overall, we found an enrichment for early HSPCs in mPB as compared to BM and observed differences in surface protein and molecular regulator expression within HSPC metaclusters, highlighting the contextual importance for defining cell states within the human.

### CB is enriched for MPPs and FL contains distinct HSPC subsets

To explore the features of adult and fetal HSPC, we performed mass cytometry on CB (n=4) and FL samples (n=3, range 17-20 PCW). As with the mPB samples, we gated CD34+ cells from BM, CB, and FL samples and performed leiden clustering and PAGA. We quantified the pairwise Manhattan distance between samples based on cell composition using leiden proportions and hierarchically clustered the resulting distance matrix (Fig. 6A). All samples clustered by tissue, and FL samples were the most different when compared to CB and BM. We observed in both fetal tissues a significant increase in the frequency of CD34+/CD38-cells when compared to BM (CB: p=0.0061, FL: p=0.017; Fig. 6B).

**Figure 6:**
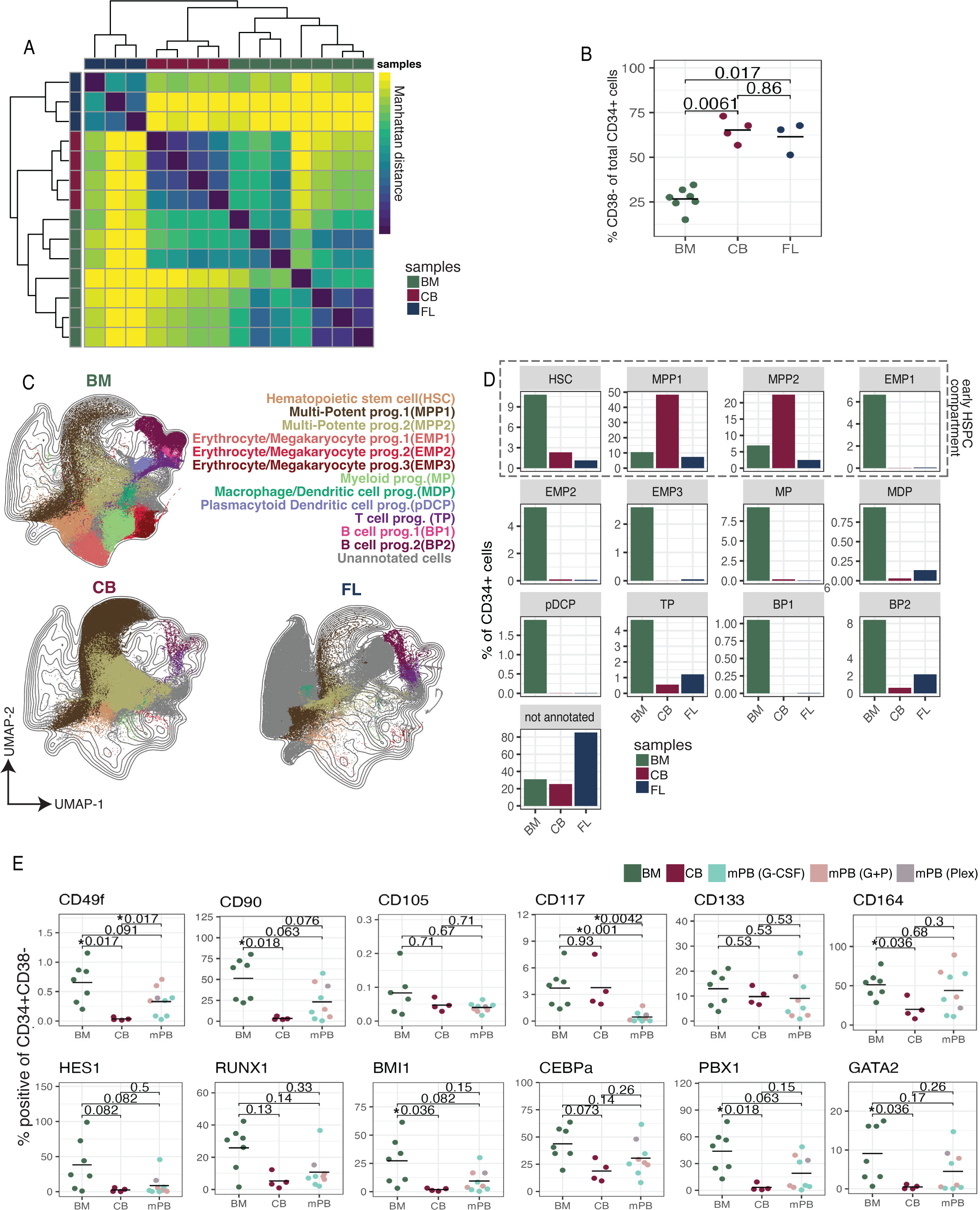
CB is enriched for MPPs and FL samples are comprised of cells with distinct expression patterns. A) Heatmap of Manhattan distance between donor samples based on leiden cluster frequencies of CD34+ cells, colored by tissue (left and top annotation colors). B) Dotplot quantifying frequency of CD38-cells within CD34+ compartment by tissue. P-value derived from Wilcoxon rank sum test. C) UMAP of BM, CB, and FL CD34+ cells colored by metacluster and separated by tissue. D) Barplots quantifying frequency of HSPC metaclusters by tissue. E) Dotplots quantifying % positivity of markers within CD34+CD38-cells, separated by tissue/product. Q-values derived from Wilcoxon rank sum test with local FDR correction.

As before, BM cells were labeled with their original BM metacluster annotations (Fig. 1), and used to train a classifier to automatically assign metacluster annotations to phenotypically similar FL and CB cells (see *Methods*). When visualized by UMAP, we observed substantial overlap between CB and BM cells, particularly MPP1 and MPP2s, in agreement with previous publication ^59^ (Fig. 6C). FL cells, however had minimal overlap with BM cells. We quantified these observations and found that over 70% of CD34+ CB cells were MPPs, while over 80% of CD34+ CB cells remained unannotated, as they had no analog in BM (Fig. 6D).

As the FL cell phenotypes and composition were substantially different from BM, we conducted a separate analysis on FL samples alone. More than 85% of the FL samples were comprised of CD34-cells (Fig. S6A), so we performed leiden clustering and PAGA on total CD45+ FL cells. Leiden clusters were metaclustered and annotated based on canonical lineage markers (*e.g.* CD3+ T cells) or descriptively based on unique cellular phenotypes (*e.g.* GATA2+Foxp1+) (Fig. S6B, Fig 3B-D). Most cells expressed at least one lineage marker, as previously reported ^60^. CD71+GATA1+ and CD71+GATA1-cells were on average, the most frequent phenotypes present in FL (Fig. S6E). While CD71 uniquely identified erythroid progenitors in our BM dataset (Fig. 2), CD71 can also be used to identify cycling cells ^61^. As the FL is a site of extensive cell proliferation ^62^ this is a plausible interpretation.

CD34+ is used clinically to identify HSPCs ^63^, and CD34+ CD38-cells enrich for the least differentiated HSPCs (Fig. 1). To survey the expression patterns of these early HSPCs across graft sources, we gated CD34+CD38-cells from BM, CB, and mPB and quantified percent positive by sample (Fig 6E). We analyzed the expression of described markers for the identification of human HSC, such as CD133, CD105, CD164, CD49f, CD117, CD90 ^8,54–57^, as well as the expression of several molecular regulators. We observed a significantly lower expression of CD49 (p=0.017), CD90 (p=0.018), CD164 (p=0.036), BMI-1 (p=0.036), PBX-1 (p=0.018), and GATA-2 (p=0.036) in CB compared to BM. These expression patterns likely reflect the enrichment for MPP phenotypes in early CB HSPCs. Early HSPCs were, however, very similar between BM and mPB, with only significantly lower expression of CD117 in mPB samples (p=0.001). In summary, we identified a substantial enrichment for MPPs in CB as compared to BM, as well as a diversity of cellular phenotypes unique to FL.

## Discussion

The self-renewal and lineage commitment fates of HSPCs is largely dictated by levels of molecular regulators and TFs. In earlier cell states with more plasticity, those components may present as promiscuous expression patterns, but eventually, become more homogenous as gene programs come under the influence of extracellular regulatory signals ^64^. The punctual transition points of hematopoiesis occur over a continuous molecular landscape, and those points can represent functionally distinct groups of cells ^14,65^. Herein, to reconcile the diversity of human CD34^+^ cells within the HSPC compartment, we leveraged mass cytometry to simultaneously capture 40+ features on millions of cells in hematopoietic tissues. These features included surface antigens for identification and prospective isolation together with a panel of molecular regulators and TFs both conserved in humans and mice. Data-driven organization revealed a molecular-regulator-centric identity across the progenitor cell spectrum while, at the same time, grounding this information within the standard model of human hematopoietic ontology that allows for prospective identification. Expression of functional markers, such as PBX1, IRF8, TdT, RUNX1, and GATA1 was critical to enhancing the separation and identification of novel putative HSPC identities.

In the classical depiction of human hematopoiesis, the segregation of lymphoid and myeloid lineages is the earliest fate decision ^66^. However, one notable observation in our study was a direct HSC to E/Meg commitment arm, the existence of which has been previously speculated. Notta *et al.* has suggested that during adult hematopoiesis, megakaryocytes can differentiate directly from very early multipotent progenitors, while still in the stem cell compartment, but not from oligopotent progenitors like CMP ^9^. In their scRNA-seq map of CD34+, Pellin *et al*. have shown that earliest fate split separates erythroid/megakaryocyte progenitors from lymphoid/myeloid progenitors ^8^. We identified three putative populations with E/Meg commitment: EMP1 that belongs to the early HSPC compartment, and EMP2 and EMP3, both of which are in the CD34+/CD38+ progenitor compartment (Fig. 1C). EMP2 and EMP3 presented a restricted expression of CD84. Indeed, in methylcellulose, CD34+CD84+ cells formed primarily erythroid colonies, whereas myeloid or mixed colonies were scarce ^67^. We also observed the expression of CD164, a sialomucin, in the EMP1 and EMP3 populations (Fig. 1C). CD164 was already described as a reliable marker for the earliest branches of HSPC specification, including its expression just beyond the earliest fate split for erythroid/megakaryocyte (E/Meg) ^8^.

To validate our approach, we prospectively isolated our putative E/Meg populations using FACS to interrogate their chromatin landscapes as well as their *in vitro* clonal growth. Our data showed that EMP1 and EMP2 cells are not fully committed to the E/Meg lineage, but in the dual Mk/E assay we see a great proportion of cells bipotent for E/Meg (Fig. 3G), supporting evidence that megakaryocytes could be derived directly from HSCs. Our functional data also confirmed the stark inflection point at the border between EMP2 and EMP3, observed in our trajectory analysis. EMP3 presented the highest accessibility of the GATA TF family and MECOM and yielded purities of >80% BUF-E across several donors and different HSPC sources (mPB and BM) in the standard MethoCult assay. EMP3 also has the highest proportion for BFU-E in the dual Mk/E assay, confirming a gradual loss of bipotentiality as cells move away from HSC in the E/Meg branch.

In human hematopoiesis, Notta *et al* also showed that previously defined myeloid progenitors are heterogeneous by comparing their lineage potential from different developmental stages ^9^. Because of an overreliance on surface markers, the standard model of hematopoiesis oversimplifies the complexity of HSPCs, yet this established paradigm has been useful for useful for understanding the differentiation process of HSC^68^. We therefore utilized both surface markers and molecular regulators to better define the differentiation trajectory of mononuclear myeloid cells and confirmed their lineage commitment with functional assays.

In the 1990’s, G-CSF-mobilized HSPCs started to become the predominant graft source for autologous transplant, due to a faster hematologic recovery. Soon after, it also became the source of the majority of allogeneic HSC transplantation, associated with improved overall patient survival ^69–71^. This change in the paradigm of clinical transplantation is due in part to the fact that mPB contains 5 to 10 times more CD34+ cells than BM aspirates and the number of HSPCs is critical for successful transplantation ^72,73^. Our data verified a significant enrichment for early HSPCs in all mPB samples, most pronounced in HSC, MPP1, and MPP2 populations when compared with steady-state adult BM (Fig. 5). We also observed the same profile in the CB samples, particularly in the MPP subpopulations (Fig. 6). CB is an alternative to BM transplantation, and it has the advantage of reduced need for HLA matching, it avails HSPC transplantation to nearly all patients ^74^. Importantly, our data produced an extensive screen of surface and molecular regulator markers of CD34+ cells in different conditions of mPB and CB samples compared with steady-state adult BM.

Our current knowledge of FL HSPC has mainly been developed in murine and *in vitro* model systems, as human fetal tissue is limited, however, a better understanding of the composition of human fetal liver and development of early hematopoiesis can lead ultimately to the production of bona fide HSC for possible clinical use ^75^. We observed that there were unique cell types/states in FL samples that were not present in the BM, likely representing fetal hematopoietic programs. While fetal and adult HSCs are known to have differences in their biology and molecular features that include cell surface markers, gene expression patterns, and cell cycle rates ^62^ our study highlights the differences across the compartment as a whole and aligns with the challenges seen in deriving equivalent adult hematopoietic cells from embryonic sources *in vitro*, like pluripotent cell systems^76^.

Altogether, we provide new insights into the earliest events of human hematopoietic immune development, new definitions for prospective isolation of HSPCs, and a reference single cell framework for further study and manipulation of these tissues in the human.

## Acknowledgements

We would like to thank Dr. Agnieszka Czechowicz, and Yan Yi (Shirley) Chan for their support with the STEMvision™ Human (Stem Cell Technologies). D.R.G. was supported by a Bio-X Stanford Interdisciplinary Graduate Fellowship, a Fred Hutchinson Cancer Center Mahan Fellowship, and a Cancer Research Institute Irvington Postdoctoral Fellowship. This work was supported by NIH grants DP2EB024246, R01AG056287, R01AG057915, R01AG068279, U19AG065156, U24CA224309, P30AG066515, U54HL165445. This study was also supported by a research grant from bluebird bio inc. who also provided much of the BM, mPB, and fetal liver studies here. Figures were created using BioRender (http://www.biorender.com) and Illustrator (Adobe).

## Author contributions

P.F. conceptualization, methodology, investigation, data curation, validation, writing-original draft preparation & editing; D.R.G. methodology, software, formal analysis, validation, data curation, writing-review & editing; L.B. conceptualization, methodology, investigation, validation; R.B. methodology, investigation, validation, software, formal analysis; W.R., D.H., and VMS: methodology, investigation, validation; D.T. software, formal analysis; I.S. resources; S.C.B. conceptualization, writing-review & editing, and led the study.

## Inclusion and diversity statement

One or more of the authors self-identifies as a member of the LGBTQ+ community and/or female ethnic minority in science.

## Resources

| REAGENT or RESOURCE | SOURCE | IDENTIFIER |
| --- | --- | --- |
| **Antibodies** |  |  |
| All antibodies used for cytometry | Various | Table S1 |
| **Cell annotations** |  |  |
| Criteria for the metaclusters | literature | Table S2 |
| **Biological Samples** |  |  |
| Human BM CD34+ cells | bluebird bio | n/a |
| Human mPB CD34+ cells | bluebird bio | n/a |
| Human MNC from FL | AllCells | allcells.com |
| **Chemicals, Peptides, and Recombinant Proteins** |  |  |
| Ficoll-Paque Plus | GE Healthcare | Cat#300-25 |
| Benzonase nuclease | Sigma-Aldrich | Cat#E1014-25KU |
| Bovine Serum Album (BSA)<br>Heat-shock Treated | Thermo Fisher<br>Scientific | Cat#BP1600-100 |
| Human TruStain FcX (Fc<br>Receptor Blocking Solution) | BioLegend | Cat#422302 |
| Cell-ID Intercalator-Ir | Standard BioTools<br>Inc. | Cat#201192A |
| Cell-ID Cisplatin | Standard BioTools<br>Inc. | Cat#201064 |
| Saponin from quillaja bark | Sigma-Aldrich | Cat#S7900-25G |
| Calibration Beads, EQ, Four<br>Element | Standard BioTools<br>Inc. | Cat#201078 |
| 7-AAD Viability Staining<br>Solution | BioLegend | Cat#420404 |
| **Deposited Data** |  |  |
| Mass cytometry batch-corrected fcs files | This study | DOI: 10.5281/zenodo.8219542 |
| Mass cytometry processed<br>data table | This study | DOI: 10.5281/zenodo.8219606 |
| ATAC-seq sequence data | This study | GEO: GSE241229 |
| ATAC-seq processed data | This study | DOI: 10.5281/zenodo.8266662 |
| **Software and Algorithms** |  |  |
| RStudio | RStudio | rstudio.com |
| MATLAB | MathWorks | mathworks.co |
| Cytobank | BeckmanCoulter | beckman.com |
| CellEngine | Primity Bio | cellengine.com |
| **Other** |  |  |
| Mass cytometry code | This study | DOI: 10.5281/zenodo.8219606 |
| ATAC-seq code | This study | DOI: 10.5281/zenodo.8266662 |

## Material and Methods

### Primary samples

All the primary samples were deindentified and purchased from suppliers of healthy donors (AllCells and StemExpress). The CD34-enriched cells (CliniMACS, Miltenyi) from BM and mPB samples from healthy donor volunteers were provided by bluebird bio. We used a total of 12 samples of CD34-enriched cells from BM, 9 samples of CD34-enriched cells from mPB (G-CSF (n=5); Plex (n=3); G-CSF+Plex (n=1)); 4 samples of CD34-enriched cells from CB, and 3 samples of mononuclear cells from FL.

### Flow cytometry screen

To design a deep phenotype antibody panel for human HSPC, we used the LEGENDScreen™ kit, which contains 342 antibodies (including 10 isotype controls), all conjugated to PE (Biolegend). CD34-enriched cells from BM samples (n=2), along with a carrier cell population (fixed Jurkat cell line) to minimize cells loss when dealing with limiting numbers of HSPC, were first stained with CD7, CD33, CD34, CD38, and 7AAD, and then stained with the kit according to the manufacturer’s instructions. Cells were analyzed using the BD LSR II Flow Cytometer (BD Biosciences). The antibody screen was performed in biological duplicate, and fixed Jurkat cells were gated out of the analysis based on the expression of 7AAD, before gating the human CD34+ cells for HSC/MPP, erythroid/megakaryocytes progenitors, lymphoid and myeloid progenitors (Fig. S1A).

### Mass cytometry antibody conjugation, staining, and data acquisition

Antibody conjugation was performed as previously described ^77^. Briefly, metal-isotope labeled antibodies used in this study were conjugated using the MaxPar X8 Antibody Labeling kit per manufacturer instruction (Standard BioTools Inc.) or were purchased from Standard BioTools Inc. pre-conjugated. Each conjugated antibody was quality checked and titrated to optimal staining concentration using a combination of primary human cells and/or cancer cell lines.

Frozen CD34-enriched cells or MNC from FL were thaw and suspended in cell staining media (CSM: PBS with 0.5% BSA, 0.02% sodium azide, and benzonase 25×108 U/mL (Sigma)) with TruStain FC blocker for 10 min at RT prior to staining. Next, the cocktail of surface staining was added (Supplemental Table1) and performed for 30 min at RT. Cells were washed in CSM and resuspended in cisplatin for 5 min to label non-viable cells (Sigma, 0.5 μM final concentration in PBS). Cells were washed in CSM, fixed with 1.6% PFA in PBS for 10 min, barcoded ^78^ and subsequently pooled to be washed with CSM once. Next, cells were permeabilized with ice cold methanol (Sigma), on ice for 10 min. Membrane permeabilized cells were washed twice with CSM, and all isotope-labeled antibodies against intracellular antigens were pre-mixed and filtered before staining in the same fashion as surface antibodies. After intracellular antibody staining, cells were washed once with CSM and then re-suspended in DNA intercalator solution (1.6% PFA diluted in PBS, 1:5000 DNA intercalator from Standard BioTools Inc.) until ready for data acquisition on the CyTOF2 mass cytometer (Standard BioTools Inc.). For the acquisition, all samples were filtered through a 35 μm nylon mesh cell strainer, resuspended at 1 × 10^6^ cells/mL in ddH2O supplemented with 1x EQ four element calibration beads (Standard BioTools Inc.). Barcoded samples were acquired using the Super Sampler injection system (Victorian Airship).

### FACS sorting

Frozen CD34-enriched cells from BM or mPB were thawed and suspended in in cell staining media (CSM: PBS with 0.5% BSA and 0.02% sodium azide and benzonase 25×108 U/mL (Sigma)) with TruStain FC blocker for 10 min at RT prior to staining. Next, the cocktail of surface antibodies was added (Table S1), and cells were stained in the dark for 30 min on ice and then washed in CSM. Prior to data acquisition, cell suspensions were spiked with 7-AAD (BioLegend) to label non-viable cells. The sorted populations (Fig. S3B, S4D) were split for CFU assay in MethoCult, CFU-Mk/E, and bulk ATAC-seq.

### CFU assay in Methylcellulose

Cells were plated in MethoCult™ H4435 Enriched (Stem Cell Technologies) per the manufacturer’s instructions. Briefly, sorted cell populations were seeded (100 or 250 cells/well) into 6 well SmartDish™ (Stem Cell Technologies). After incubation for 14 days, at 37°C in 5% CO_2_, hematopoietic colony-forming units were automated counted, and analyzed by STEMvision™ Human (Stem Cell Technologies).

### Dual differentiation assay

Dual Mk/E colony assay was performed as described before ^46^. Briefly, sorted 250 cells/well were plated in a 6 well plate of MegaCult C Medium plus Lipids (Stem Cell Technologies) with rhEPO (recombinant human erythropoietin), recombinant human interleukin rhIL-3, rhIL-6, and rhSCF (recombinant human stem cell factor), rhTPO (recombinant human thrombopoietin). All cytokines were purchased from Stem Cell Technologies. After 2 weeks colonies were stained with the following antibodies: CD71/CD235a-PE and CD41a-FITC. Plates were read on Image Xpress Micro XLS (Molecular Devices), and colonies were scored based on the fluorescence signal as megakaryocyte only (CFU-Mk), erythroid only burst forming unit (BFU)-E, or megakaryocyte + erythroid (CFU-Mk/E).

### Mass cytometry data pre-processing

Fcs files from the mass cytometer were bead normalized ^79^ and debarcoded using the R package, *premessa*. Files were uploaded to Cytobank and leukocytes were gated as DNA^+^ bead^-^ viability^-^ and CD45^+^ (Fig. S2C-D). Fcs files were downloaded and compensated with the R package, *CATALYST* ^80^. Files were transformed with hyperbolic arcsin function using a cofactor of 5.

Batch correction was performed utilizing a common bone marrow sample included with each batch as previously described^81^. For each protein channel, the median expression value was determined for all common samples. The difference in medians was subtracted from all events to shift the distributions downward to have identical medians. Negative values that were introduced by this process were reset to the absolute value of a random normal distribution centered at zero to align with the noise floor for each channel. The resulting distributions were then 99.8^th^ percentile normalized downward. The transforms applied to each batch common sample were then applied to all samples from that batch. This approach retained biological heterogeneity while accommodating batch correction for a range of distributions and batch effect manifestations.

### Gating, clustering, and pseudotime analysis

Batch-corrected fcs files were uploaded to CellEngine and gated into lin-CD34+, CD34-, and HSPC populations (Fig. 1B). Files were downloaded and analyzed in R. All mass cytometry analysis was performed on asinh-transformed (cofactor 5) data, with each protein channel scaled to the 99.9^th^ percentile. Samples were equally downsampled by tissue and donor to a total of 400k total cells (BM analysis; FL analysis), 400k CD34+ cells (BM and mPB analysis) or 300k CD34+ cells (BM, CB, and FL analysis).

PAGA ^25^ from SCANPY ^82^ was called from R using the reticulate package. The pipeline was individually performed for each analysis, including additionally for BM erythroid and myeloid trajectory inference. All proteins were used for all analyses, except erythroid and myeloid analyses, which only used the subset of proteins expressed by those cells. Briefly, a knn graph was made for all cells and then used for dimensionality reduction by diffusion maps ^83^. A second knn graph was made based on these diffusion coordinates and used for Leiden clustering ^24^. A PAGA graph was inferred from the Leiden clusters and used to initialize UMAP dimensionality reduction ^28^, which was performed also using the second knn graph. Pseudotime based on the PAGA graph was calculated starting from a randomly selected HSC. For pseudotime heatmaps, cells were divided into 200 equally sized bins.

For each analysis, cells were over-clustered to generate many more Leiden clusters than expected unique cell populations. This conservative strategy avoids spurious merging of small populations into large populations and enables downstream merging and annotation, as others have noted^84^. BM and FL clusters were manually annotated into metaclusters based on surface and regulatory factor expression profiles. For analysis of mPB and CB, new Leiden clusters were derived, but BM cells retained their original metacluster annotations from the BM-only analysis. Those labels were used to train a knn-classifier to annotate cells derived from other tissues, stipulating that only BM cells within the same Leiden cluster can be considered in the knn graph. This facilitated automatic annotation of previously defined HSPCs, while preventing spurious annotation of cells with unique phenotypes not found in BM.

For erythroid and myeloid trajectory analyses (Fig. 2, 4), only the subset of BM cells in that trajectory were included in the analysis. Cells in each trajectory were selected based on their Leiden cluster and metacluster assignments (Fig. 1). The erythroid trajectory included only BM clusters connected by the PAGA graph and annotated as HSC, EMP1-3, immature erythroid, or unannotated. The myeloid trajectory included only BM clusters connected by the PAGA graph and annotated as HSC, MPP2, MP, MDP, pDCP, pDC, Monocyte, or unannotated.

### Sample Dissimilarity Metric

To calculate sample (and tissue) dissimilarity where indicated, the pairwise Manhattan distance between samples were calculated based on the vector of Leiden clusters proportions for mass cytometry data from each sample. This unbiased approach does not rely on identifying the optimal number of clusters nor the correct user annotations. Because the data are over-clustered, each true cell population is comprised of multiple Leiden clusters with small differences in protein expression profiles. This distance metric is therefore sensitive to both large changes in overall cell composition as well as smaller changes in protein expression within cell populations.

### Statistical analyses

All pairwise comparisons between samples employed the Wilcoxon rank sum test. Local FDR for multiple hypothesis correction was applied where appropriate.

### Fast-ATACseq

Cells were processed as described previously ^36^. Briefly, ∼50 000 FACS-purified cells were pelleted and used for the transposition reaction of each technical replicate, followed by DNA purification. Amplification and purification of the transposed fragments were done as described previously ^85^ with modified primers ^86^. Libraries were quantified using qPCR. All Fast-ATAC libraries were sequenced on an Illumina NovaSeq through Novogene. There were two technical replicates and two biological replicates for each population analyzed here.

### ATAC seq processing

All fastq files from Novogene were equally subsampled to 15M reads before processing using the pepATAC pipeline ^87^. Within the pipeline, reads were trimmed, and reads mapping to mitochondrial reads were filtered before aligning to the hg19 genome using Bowtie2^88^. Deduplicated reads were removed using Samblaster, and bam files were sorted and indexed using Samtools ^89^. The MACS2 algorithm was used for calling 500bp peaks (extended 250bp in either direction from the peak summit) with parameters (--shift-75--extsize 150 --nomodel --call-summits --nolambda --keep-dup all -p 0.01) ^90^. The resulting bam and bed files were used for downstream analysis.

### ATAC seq analysis

Bam and bed files were read into the ChrAccR R package to create a bulkATAC object^91^. QC metrics and plots of TSS scores and fragment length distributions were created using in-built functions in ChrAccR (scripts included in Zenodo:8266662). BulkATAC objects were then simulated as single cells before projecting in UMAP space of previously published scATAC data ^92^. The scATAC data was processed using the ArchR package, and projection was carried out using the projectBulkATAC function ^44,93^. The closest single cells to projected cell centroids by Mahalanobis distance were labeled with bulkATAC sample names. We then added motif annotations to all single cells in the dataset using the ArchR function addMotifAnnotations with the cisbp TF database ^94^. To find significantly accessible motifs in a scATAC population corresponding to a bulkATAC sample, we first found enriched peaks in the population with a Wilcoxon pairwise test (cutoff of FDR <= 0.05 & Log2FC >= 2) between the population-of-interest and all other populations as background using the getMarkerFeatures function from ArchR. Finally, we ran the peakAnnoEnrichment function on differential peaks and plotted rank-sorted TF motifs in the highly accessible peaks within the sample population by significance of negative log10 adjusted P value.

