## supplemental figures and tables for "Unravelling human hematopoietic progenitor cell diversity through association with intrinsic regulatory factors"

**Figure S1**

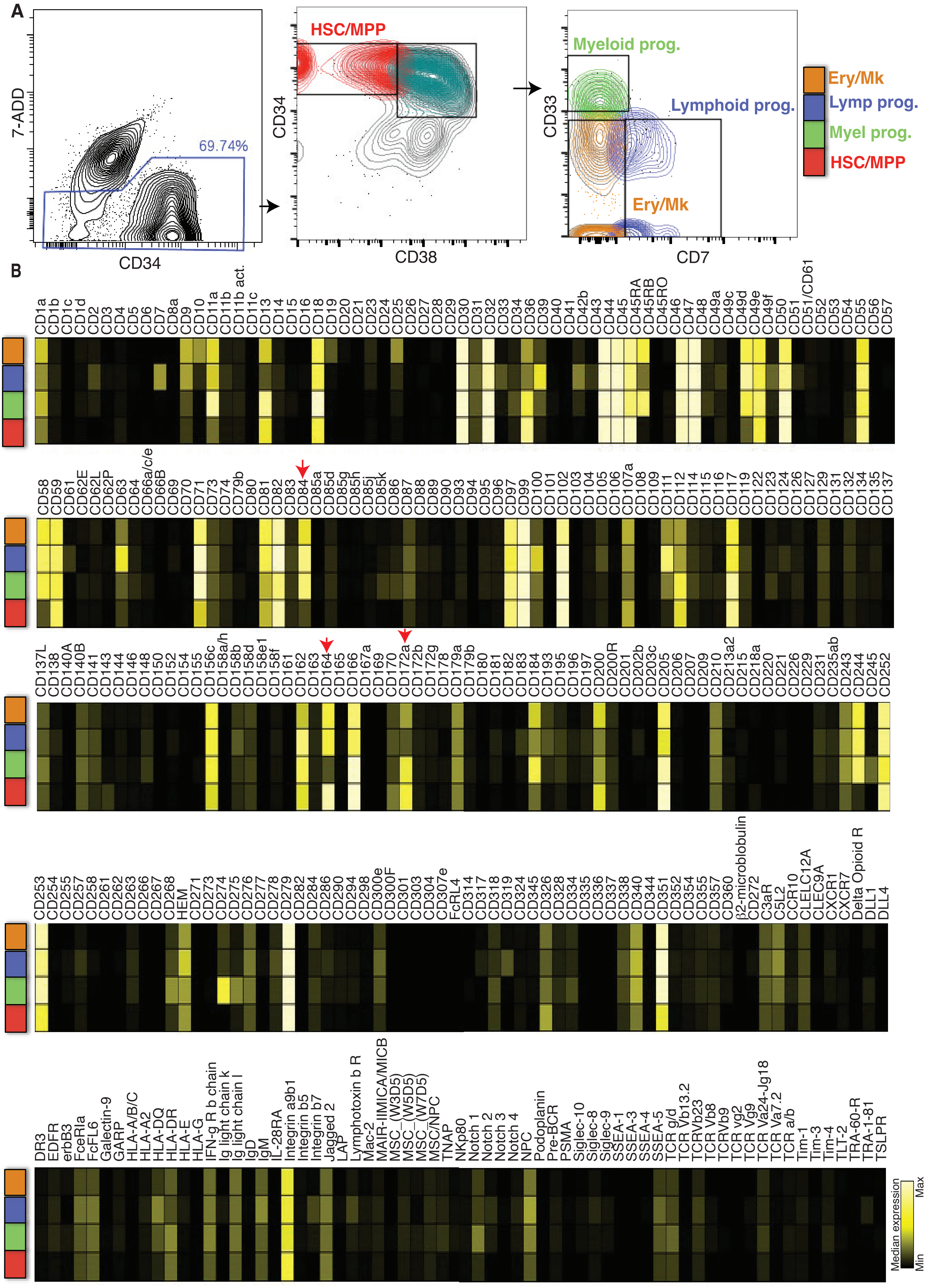
A
7-ADD
CD34
69.74%
HSC/MPP
CD34
CD38
Myeloid prog.
Lymphoid prog.
Ery/Mk
CD33
CD7
Ery/Mk
Lymp prog.
Myel prog.
HSC/MPP
B
CD1a CD1b CD1c CD1d CD2 CD3 CD4 CD5 CD6 CD7 CD8a CD9 CD10 CD11a CD11b CD11b act. CD11c CD13 CD14 CD15 CD16 CD18 CD19 CD20 CD21 CD23 CD24 CD25 CD26 CD27 CD28 CD29 CD30 CD31 CD32 CD33 CD34 CD36 CD39 CD40 CD41 CD42b CD43 CD44 CD45 CD45RA CD45RB CD45RO CD46 CD47 CD48 CD49a CD49c CD49d CD49e CD49f CD50 CD51 CD51/CD61 CD52 CD53 CD54 CD55 CD56 CD57
CD58 CD59 CD61 CD62E CD62L CD62P CD63 CD64 CD66a/c/e CD66B CD69 CD70 CD71 CD73 CD74 CD79b CD80 CD81 CD82 CD83 CD84 CD85a CD85d CD85g CD85h CD85j CD85k CD86 CD87 CD88 CD89 CD90 CD93 CD94 CD95 CD96 CD97 CD99 CD100 CD101 CD102 CD103 CD104 CD105 CD106 CD107a CD108 CD109 CD111 CD112 CD114 CD115 CD116 CD117 CD119 CD122 CD123 CD124 CD126 CD127 CD129 CD131 CD132 CD134 CD135 CD137
CD137L CD138 CD140A CD140B CD141 CD143 CD144 CD146 CD148 CD150 CD152 CD154 CD155 CD156c CD158a/h CD158b CD158d CD158e1 CD158f CD161 CD162 CD163 CD164 CD165 CD166 CD167a CD169 CD170 CD172a CD172b CD172g CD178 CD179a CD179b CD180 CD181 CD182 CD183 CD184 CD193 CD195 CD196 CD197 CD200 CD200R CD201 CD202b CD203c CD205 CD206 CD207 CD209 CD210 CD213a2 CD215 CD218a CD220 CD221 CD226 CD229 CD231 CD235ab CD243 CD244 CD245 CD252
CD253 CD254 CD255 CD257 CD258 CD261 CD262 CD263 CD266 CD267 CD268 HEM CD271 CD273 CD274 CD275 CD276 CD277 CD278 CD279 CD282 CD284 CD286 CD290 CD294 CD298 CD300e CD300F CD301 CD303 CD304 CD307e FcRL4 CD314 CD317 CD318 CD319 CD324 CD345 CD326 CD328 CD334 CD335 CD336 CD337 CD338 CD340 CD344 CD351 CD352 CD354 CD355 CD357 CD360 β2-microblobulin CD272 C3aR C5L2 CCR10 CLELC12A CLEC9A CXCR1 CXCR7 Delta Opioid R DLL1 DLL4
DR3 EDFR erbB3 FceRIa FcFL6 Galectin-9 GARP HLA-A/B/C HLA-A2 HLA-DQ HLA-DR HLA-E HLA-G IFN-g R b chain Ig light chain k Ig light chain l IgD IgM IL-28RA Integrin a9b1 Integrin b5 Integrin b7 Jagged 2 LAP Lymphotoxin b R Mac-2 MAIR-IIMICA/MICB MSC_(W3D5) MSC_(W5D5) MSC_(W7D5) MSC/NPC TNAP NKp80 Notch 1 Notch 2 Notch 3 Notch 4 NPC Podoplanin Pre-BCR PSMA Siglec-10 Siglec-8 Siglec-9 SSEA-1 SSEA-3 SSEA-4 SSEA-5 TCR g/d TCR Vb13.2 TCRVb23 TCR Vb8 TCRVb9 TCR vg2 TCR Vg9 TCR Va24-Jg18 TCR Va7.2 TCR a/b Tim-1 Tim-3 Tim-4 TLT-2 TRA-60-R TRA-1-81 TSLPR
Median expression
Max
Min

**Figure S2**

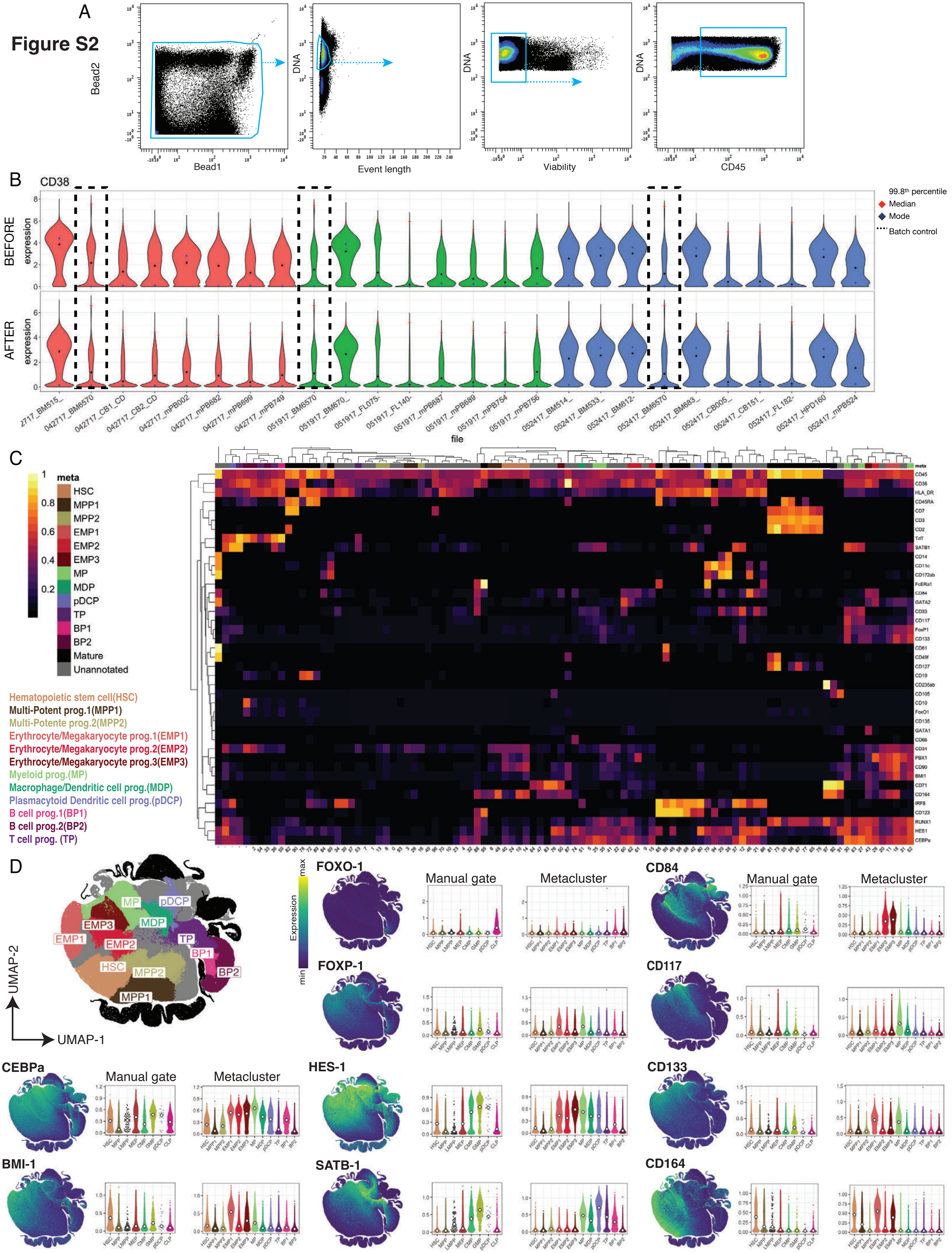

A
Bead2
Bead1
DNA
Event length
Viability
CD45
B
CD38
BEFORE
AFTER
expression
file
99.8th percentile
Median
Mode
Batch control
C
meta
HSC
MPP1
MPP2
EMP1
EMP2
EMP3
MP
MDP
pDCP
TP
BP1
BP2
Mature
Unannotated
Hematopoietic stem cell(HSC)
Multi-Potent prog.1(MPP1)
Multi-Potente prog.2(MPP2)
Erythrocyte/Megakaryocyte prog.1(EMP1)
Erythrocyte/Megakaryocyte prog.2(EMP2)
Erythrocyte/Megakaryocyte prog.3(EMP3)
Myeloid prog.(MP)
Macrophage/Dendritic cell prog.(MDP)
Plasmacytoid Dendritic cell prog.(pDCP)
B cell prog.1(BP1)
B cell prog.2(BP2)
T cell prog. (TP)
D
UMAP-1
UMAP-2
Expression
max
min
FOXO-1
FOXP-1
CD84
CD117
CEBPa
HES-1
CD133
BMI-1
SATB-1
CD164
Manual gate
Metacluster

### Figure S3

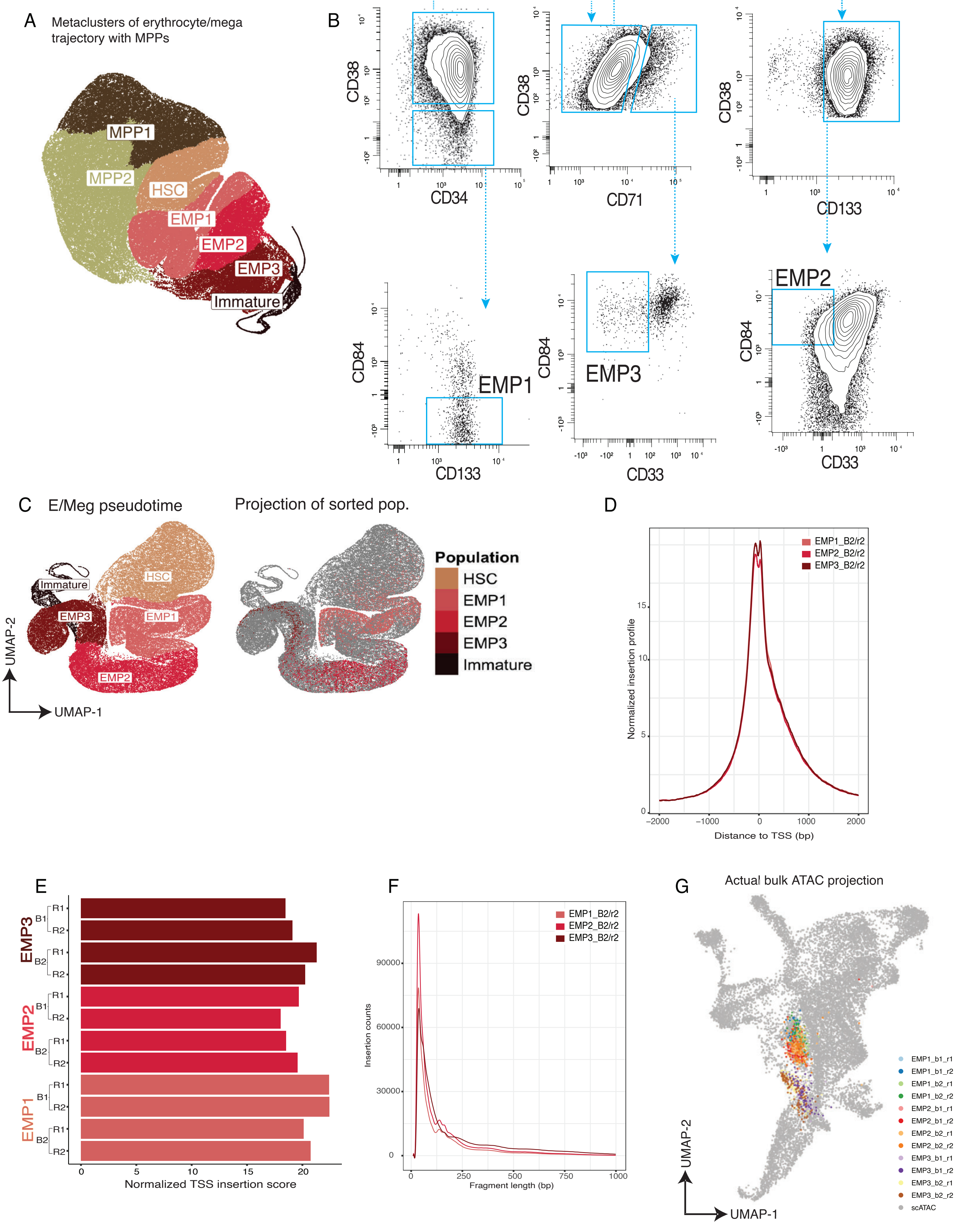

A
Metaclusters of erythrocyte/mega trajectory with MPPs
MPP1
MPP2
HSC
EMP1
EMP2
EMP3
Immature
B
CD38
CD34
CD71
CD133
CD84
CD33
EMP1
EMP2
EMP3
C
E/Meg pseudotime
Projection of sorted pop.
UMAP-2
UMAP-1
Population
HSC
EMP1
EMP2
EMP3
Immature
D
EMP1_B2/r2
EMP2_B2/r2
EMP3_B2/r2
Normalized insertion profile
Distance to TSS (bp)
E
EMP3
EMP2
EMP1
Normalized TSS insertion score
F
Insertion counts
Fragment length (bp)
G
Actual bulk ATAC projection
EMP1_b1_r1
EMP1_b1_r2
EMP1_b2_r1
EMP1_b2_r2
EMP2_b1_r1
EMP2_b1_r2
EMP2_b2_r1
EMP2_b2_r2
EMP3_b1_r1
EMP3_b1_r2
EMP3_b2_r1
EMP3_b2_r2
scATAC

Figure S4

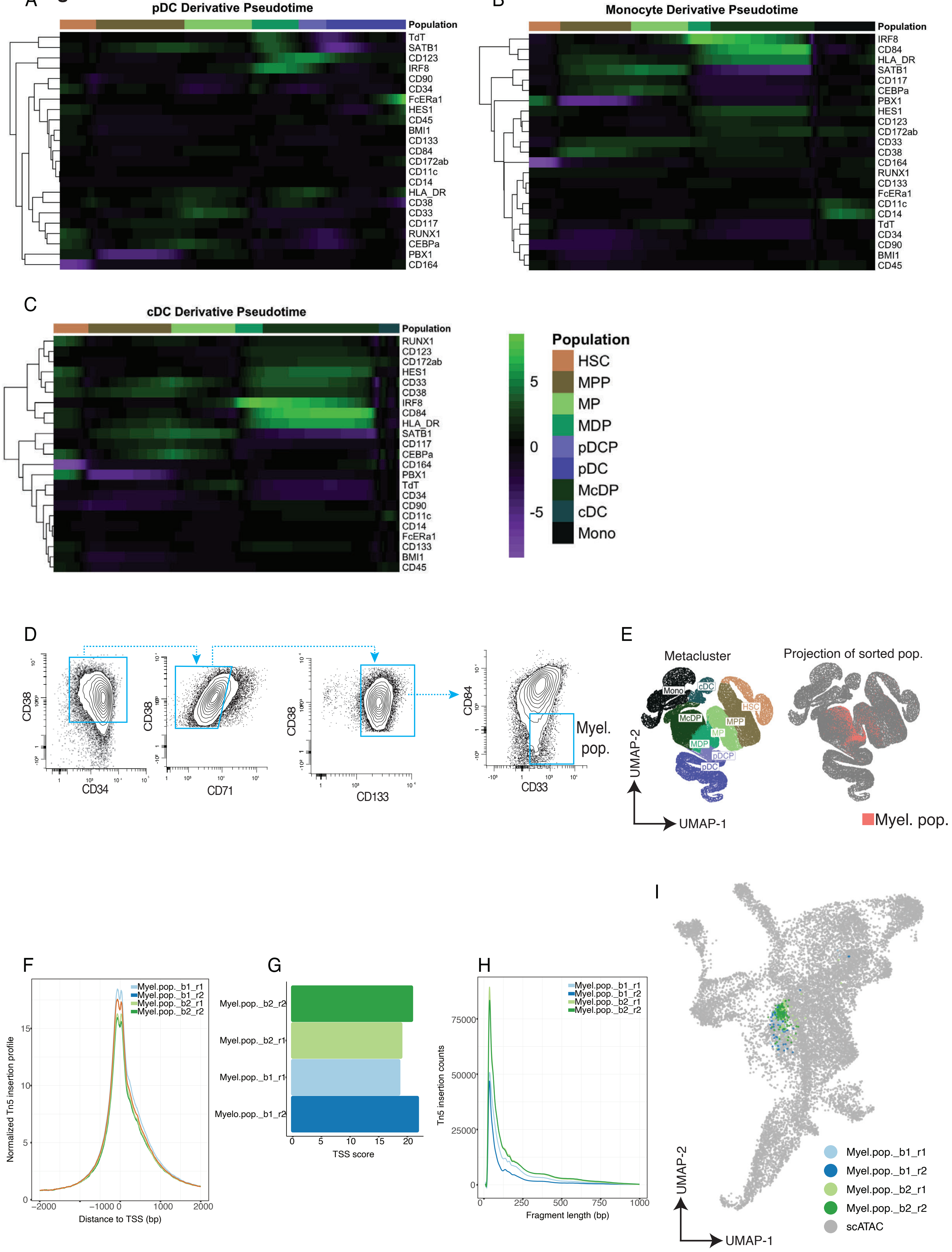
A
pDC Derivative Pseudotime
Population
TdT
SATB1
CD123
IRF8
CD90
CD34
FcERa1
HES1
CD45
BMI1
CD133
CD84
CD172ab
CD11c
CD14
HLA_DR
CD38
CD33
CD117
RUNX1
CEBPa
PBX1
CD164
B
Monocyte Derivative Pseudotime
Population
IRF8
CD84
HLA_DR
SATB1
CD117
CEBPa
PBX1
HES1
CD123
CD172ab
CD33
CD38
CD164
RUNX1
CD133
FcERa1
CD11c
CD14
TdT
CD34
CD90
BMI1
CD45
C
cDC Derivative Pseudotime
Population
RUNX1
CD123
CD172ab
HES1
CD33
CD38
IRF8
CD84
HLA_DR
SATB1
CD117
CEBPa
CD164
PBX1
TdT
CD34
CD90
CD11c
CD14
FcERa1
CD133
BMI1
CD45
5
0
-5
Population
HSC
MPP
MP
MDP
pDCP
pDC
McDP
cDC
Mono
D
CD38
CD34
CD38
CD71
CD38
CD133
CD84
CD33
Myel.
pop.
E
Metacluster
Projection of sorted pop.
Mono
cDC
HSC
McDP
MPP
MP
MDP
pDCP
pDC
UMAP-2
UMAP-1
Myel. pop.
F
Myel.pop._b1_r1
Myel.pop._b1_r2
Myel.pop._b2_r1
Myel.pop._b2_r2
Normalized Tn5 insertion profile
Distance to TSS (bp)
G
Myel.pop._b2_r2
Myel.pop._b2_r1
Myel.pop._b1_r1
Myelo.pop._b1_r2
TSS score
H
Myel.pop._b1_r1
Myel.pop._b1_r2
Myel.pop._b2_r1
Myel.pop._b2_r2
Tn5 insertion counts
Fragment length (bp)
I
Myel.pop._b1_r1
Myel.pop._b1_r2
Myel.pop._b2_r1
Myel.pop._b2_r2
scATAC
UMAP-2
UMAP-1

### Figure S5

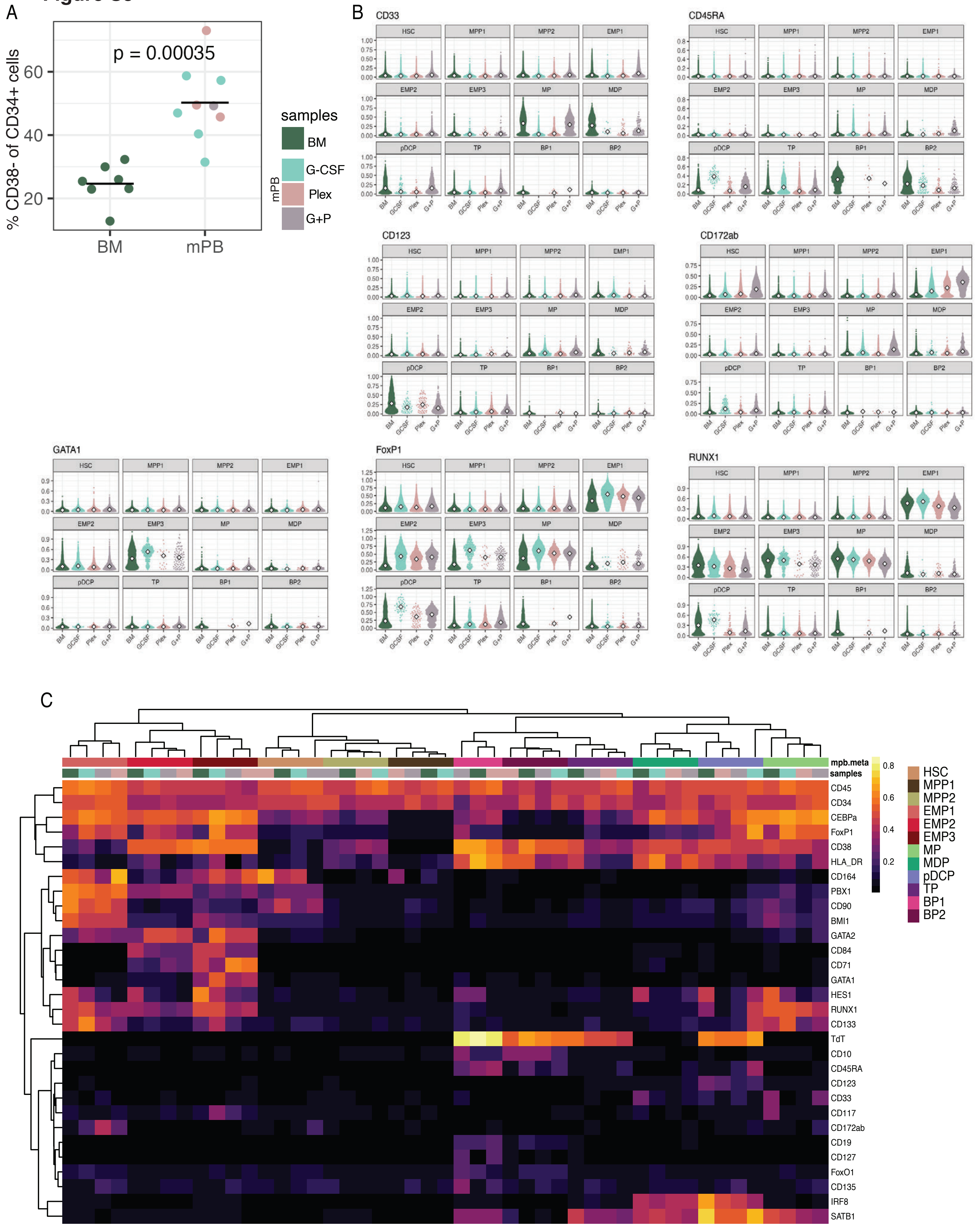
A
B
C
p = 0.00035
% CD38- of CD34+ cells
BM
mPB
samples
BM
G-CSF
Plex
G+P
CD33
CD45RA
CD123
CD172ab
GATA1
FoxP1
RUNX1
HSC
MPP1
MPP2
EMP1
EMP2
EMP3
MP
MDP
pDCP
TP
BP1
BP2
GCSF
mpb.meta
samples
CD45
CD34
CEBPa
FoxP1
CD38
HLA_DR
CD164
PBX1
CD90
BMI1
GATA2
CD84
CD71
GATA1
HES1
RUNX1
CD133
TdT
CD10
CD45RA
CD123
CD33
CD117
CD172ab
CD19
CD127
FoxO1
CD135
IRF8
SATB1

**Figure S6**

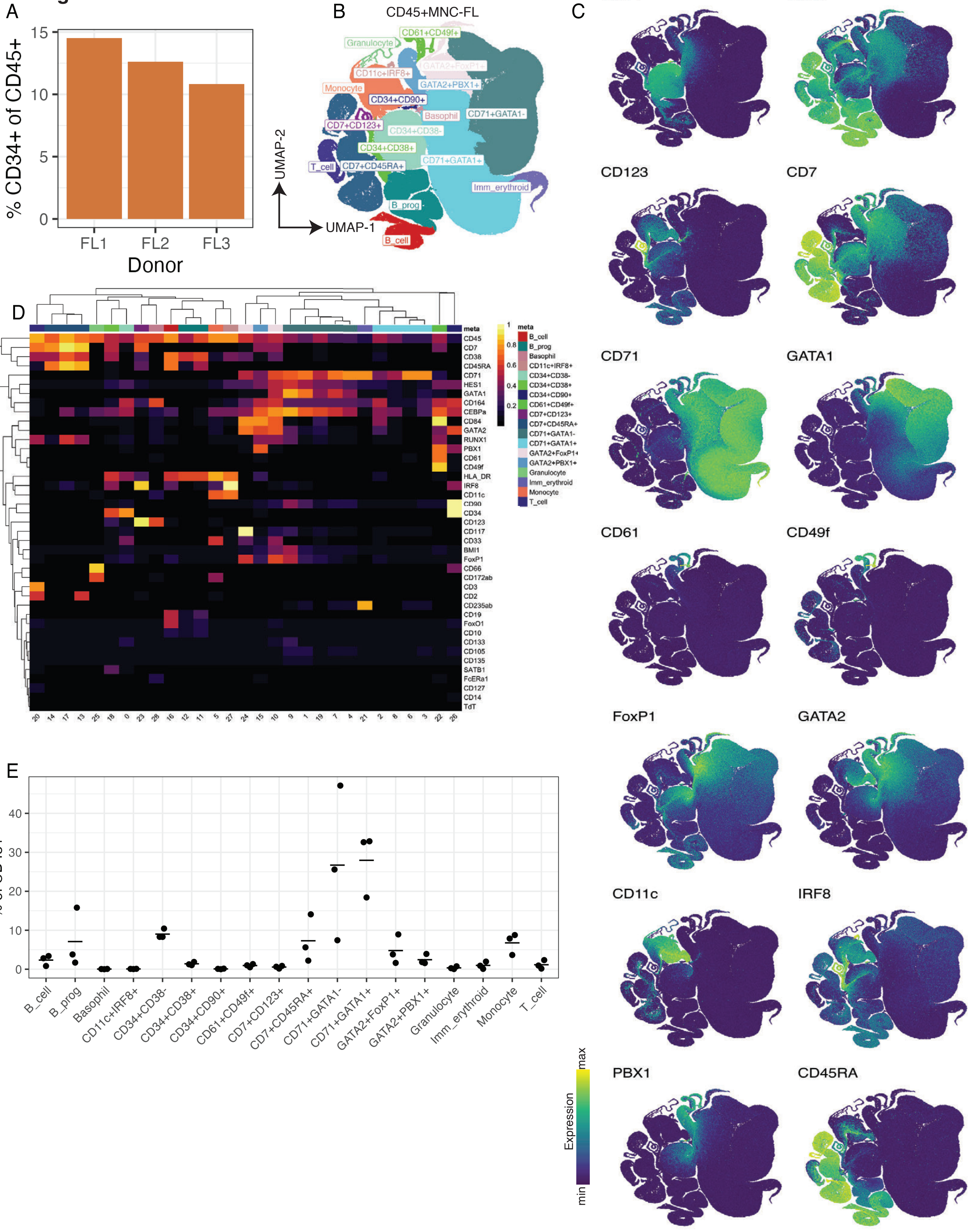
A
% CD34+ of CD45+
15
10
5
0
FL1
FL2
FL3
Donor
B
CD45+MNC-FL
Granulocyte
CD61+CD49f+
GATA2+FoxP1+
CD11c+IRF8+
GATA2+PBX1+
Monocyte
CD34+CD90+
Basophil
CD71+GATA1-
CD7+CD123+
CD34+CD38-
CD34+CD38+
T_cell
CD7+CD45RA+
CD71+GATA1+
Imm_erythroid
B_prog
B_cell
UMAP-2
UMAP-1
C
CD34
CD38
CD123
CD7
CD71
GATA1
CD61
CD49f
FoxP1
GATA2
CD11c
IRF8
PBX1
CD45RA
Expression
max
min
D
meta
CD45
CD7
CD38
CD45RA
CD71
HES1
GATA1
CD164
CEBPa
CD84
GATA2
RUNX1
PBX1
CD61
CD49f
HLA_DR
IRF8
CD11c
CD90
CD34
CD123
CD117
CD33
BMI1
FoxP1
CD66
CD172ab
CD3
CD2
CD235ab
CD19
FoxO1
CD10
CD133
CD105
CD135
SATB1
FcERa1
CD127
CD14
TdT
20 14 17 13 25 18 0 23 28 16 12 11 5 27 24 15 10 9 1 19 7 4 21 2 8 6 3 22 26
1
0.8
0.6
0.4
0.2
meta
B_cell
B_prog
Basophil
CD11c+IRF8+
CD34+CD38-
CD34+CD38+
CD34+CD90+
CD61+CD49f+
CD7+CD123+
CD7+CD45RA+
CD71+GATA1-
CD71+GATA1+
GATA2+FoxP1+
GATA2+PBX1+
Granulocyte
Imm_erythroid
Monocyte
T_cell
E
% of CD45+
40
30
20
10
0
B_cell
B_prog
Basophil
CD11c+IRF8+
CD34+CD38-
CD34+CD38+
CD34+CD90+
CD61+CD49f+
CD7+CD123+
CD7+CD45RA+
CD71+GATA1-
CD71+GATA1+
GATA2+FoxP1+
GATA2+PBX1+
Granulocyte
Imm_erythroid
Monocyte
T_cell

**Supplementary Table 1: Mass and Flow Cytometry Panels**

| HSPC mass cytometry panel | | | | | |
|---|---|---|---|---|---|
| Antibody target | Clone | Provider | Mass channel | Isotope | Staining concentration (mg/mL) |
| CD45 | HI30 | Fluidigm | 89 | Y | 1 |
| CD235ab | HIR2 | Biolegend | 113 | In | 1 |
| CD71 | CY1G4 | Biolegend | 115 | In | 0.5 |
| CD61 | VI-PL2 | Biolegend | 139 | La | 0.5 |
| CD3 | UCHT1 | Biolegend | 141 | Pr | 0.5 |
| CD19 | HIB19 | Fluidigm | 142 | Nd | 2 |
| CD90-Biotin | 5E10 | BD | 143 | Nd | 5 |
| CD14 | M5E2 | Biolegend | 144 | Nd | 4 |
| Hes1 | 3A3 | Abnova | 145 | Nd | 0.75 |
| CD164 | 67D2 | Biolegend | 146 | Nd | 3 |
| CD42b | HIP1 | Biolegend | 147 | Sm | 4 |
| CD34 | 581 | Biolegend | 148 | Nd | 1 |
| IRF-8 | 7G11A45 | Biolegend | 149 | Sm | 2 |
| CD105 | 43A3 | Biolegend | 150 | Nd | 4 |
| CD123 | 6H6 | Fluidigm | 151 | Eu | 1 |
| CD10 | HI10a | Biolegend | 152 | Sm | 1 |
| FcER1 | AER-37 | Biolegend | 153 | Eu | 2 |
| CD84 | CD84.1.21 | Fluidigm | 154 | Sm | 1 |
| CD110 | 1.6.1 | BD | 155 | Gd | 2 |
| PBX1 | polyclonal | CST | 156 | Gd | 1 |
| Runx1 | 1C5B16 | Biolegend | 157 | Gd | 3 |
| CD33 | WM53 | fluidigm | 158 | Gd | 1 |
| CD11c | 3.9 | fluidigm | 159 | Td | 1 |
| GATA-1 | D52H6 | CST | 160 | Gd | 3 |
| C/EBPa | 16C12B70 | Biolegend | 161 | Dy | 2 |
| CD41 | HIP8 | Biolegend | 162 | Dy | 0.5 |
| CD36 | 5-271 | Biolegend | 163 | Dy | 0.5 |
| CD49f | GoH3 | Biolegend | 164 | Dy | 3 |
| CD127 | A019D5 | Fluidigm | 165 | Ho | 1 |
| FoxP1 | D35D10 | CST | 166 | Er | 2 |
| CD66 | B1.1 | BD | 167 | Er | 1 |
| CD38 | HIT2 | Biolegend | 168 | Er | 2 |
| CD45RA | HI100 | Fluidigm | 169 | Tm | 1 |
| CD135-APC | BV10A44H2 | Biolegend | 170 | Er | 5 ul |
| CD44 | IM7 | Fluidigm | 171 | Yb | 1 |
| CD133-PE | AC133 | MT | 172 | Yb | 8 ul |
| CD172ab | SE5A5 | Biolegend | 173 | Yb | 3 |
| CD2 | TS1 | Biolegend | 174 | Yb | 1 |

| | | | | | |
|---|---|---|---|---|---|
| GATA-2 | 2D11 | Abnova | 175 | Lu | 0.5 |
| SATB1 | SAT-5 | Sigma-Aldrich | 176 | Yb | 3 |
| HLA-DR | L243 | Biolegend | 209 | Bi | 1 |

**novel EMP and MP FACS Panel**

| Antibody target | Clone | Provider | Fluorochrome | Isotope | Staining concentration (mg/mL) |
|---|---|---|---|---|---|
| CD34 | AC136 | MT | VioBlue | IgG2a | 2 |
| CD38 | HIT2 | Biolegend | APC-Cy7 | IgG1 | 2 |
| CD33 | WM53 | Biolegend | PE-Cy7 | IgG1 | 0.2 |
| CD71 | CY1G4 | Biolegend | PE | IgG2a | 0.5 |
| CD84 | CD84.1.21 | Biolegend | APC-Cy7 | IgG2a | 1.2 |
| CD133 | AC133 | MT | VioBright 515 | IgG1 | 1 |

**Conventional MEP FACS Panel**

| Antibody target | Clone | Provider | Fluorochrome | Isotope | Staining concentration (mg/mL) |
|---|---|---|---|---|---|
| CD34 | AC136 | MT | FITC | IgG2a | 1.5 |
| CD38 | HIT2 | Biolegend | BV421 | IgG1 | 0.2 |
| CD45RA | HI100 | Biolegend | AF700 | IgG2b | 2 |
| CD10 | HI10a | BD | BV650 | IgG1 | 2 |
| CD123 | 6H6 | Biolegend | PE-Cy7 | IgG1 | 0.6 |

**Supplementary Table 2: Criteria for annotating the metacluster within CD34+ cells and the frequency of CD34+ cells in each tissue**

| | | % of CD34+ (mean ± sd) | | | | |
|---|---|---|---|---|---|---|
| **Population** | **Markers** | **Bone Marrow** | **Mobilized Peripheral Blood** | **Cord Blood** | **Fetal Liver** | **References** |
| BP1 | CD38+/FOXP1+/CD19-/ TDT+ | 1.1 ± 0.41 | 0.016 ± 0.021 | 0 | 0 | 1,2 |
| BP2 | CD38+/CD19+/TDT+ | 8.4 ± 8.6 | 0.063 ± 0.062 | 0.79 ± 0.67 | 1.9 ± 0.99 | 2 |
| EMP1 | CD38-/GATA2+/FOXP1+ | 6.2 ± 5.6 | 2.9 ± 6 | 0.029 ± 0.036 | 0.055 ± 0.024 | 3,4,5,6 |
| EMP2 | CD38+/CD71mid/GATA1+/GATA2+ | 5.6 ± 2.8 | 1.1 ± 1 | 0.091 ± 0.092 | 0.069 ± 0.014 | 3,4,5,6 |
| EMP3 | CD38+/CD71high/GATA1+ | 2.5 ± 2.6 | 0.21 ± 0.41 | 0 | 0.051 ± 0.026 | 3,4,5,6 |
| HSC | CD38-/PBX1+/CD90+ | 11 ± 4.5 | 16 ± 13 | 2.9 ± 2.6 | 1 ± 0.89 | 7 |
| MDP | CD38+/CD33+/IRF8+/TDT-/HLADR+ | 0.93 ± 0.32 | 0.043 ± 0.035 | 0.025 ± 0.025 | 0.13 ± 0.015 | 8 |
| MP | CD38+CD33+/HLA-DR+ | 9.2 ± 6.5 | 1.5 ± 2.5 | 0.33 ± 0.43 | 0.051 ± 0.03 | 9 |
| MPP1 | CD38-/CD90- | 11 ± 8.4 | 42 ± 17 | 50 ± 3.2 | 6.5 ± 8 | 9 |
| MPP2 | CD38 med/GATA2-/PBX1- | 7.1 ± 4.5 | 16 ± 7.6 | 20 ± 4.8 | 2.2 ± 2.4 | |
| TP | CD38+/SATB1+/TDT+/CD19- | 4.7 ± 1.8 | 1 ± 0.66 | 0.61 ± 0.43 | 1 ± 1.2 | 10 |
| pDCP | CD38+/TDT+/IRF8+/CD123high | 1.9 ± 0.92 | 0.14 ± 0.18 | 0.0063 ± 0.0053 | 0.016 ± 0.018 | 11 |

Pop.: population; BM: bone marrow; mPB: mobilized peripheral blood; CB: cord blood; FL: fetal liver; BP1: B cell progenitor ; BP1: B cell progenitor 2; EMP1: erythrocyte/megakaryocyte progenitor 1; EMP2: erythrocyte/megakaryocyte progenitor 2; EMP3: erythrocyte/megakaryocyte progenitor 3; HSC: hematopoietic stem cell; MDP: macrophage/dendritic cell progenitor; MPP1: multi-potent progenitor 1; MPP2: multi-potent progenitor 2; TP: T cell progenitor; pDCP: plasmocytoid dendritic cell progenitor.

**Supplemental Information**

**Figure S1: Human cell surface molecule flow cytometry screen– related to Fig. 1.**

A) Gating strategy for HSPC populations. B) Median expression of molecules in the screen separated by HSPC populations. Sidebar colors same as in A). Arrows indicate markers called out in text.

**Figure S2: Data preprocessing, leiden clustering and marker expression – related to Fig. 1.**

A) Gating strategy for viable CD45+ cells. B) Exemplative violin plots of CD38 expression of all mass cytometry data used in the study, separated by donor/tissue and colored by experimental batch before (top) and after (bottom) batch correction. Batch control samples used to calculate batch corrected boxed by dashed lines. C) Heatmap of median expression of leiden clusters, colored by metacluster annotation (top). D) UMAPs colored by protein expression (left) alongside violin plots of the same protein separated by manual gate (center) and metacluster (right) as in Fig. 1F. Diamond indicates median.

**Figure S3: EMP populations sorting strategy and ATAC-seq quality control – related to Fig. 3.**

A) UMAP of BM cells in the erythroid trajectory with the addition of the MPPs clusters to the analysis. B) FACS strategy for EMP populations used in ATAC-seq and *in vitro* clonal differentiation assays. C) UMAP of BM cells in the erythroid trajectory, colored by metacluster (left) or projected FACS populations (right). D) Histogram of normalized Tn5 insertion profiles centered at transcription start sites (TSSs) for the indicated ATAC-seq libraries. E) Barplots of normalized TSS insertion scores across biological and technical replicates of cell populations. F) Histogram of insertion counts by fragment length for indicated ATAC libraries. G) UMAP previously published and annotated BM scATAC dataset with the projection of biological and technical replicates of ATAC-seq libraries.

**Figure S4: Myeloid trajectories, sorting strategy, and ATAC-seq quality control – related to Fig. 4.**

A) Heatmap of first derivative of marker expression, showing rate of expression change, along the pDC pseudotime trajectory. B) Same as A) for monocyte trajectory. C) Same as A) for cDC trajectory. D) FACS strategy for myeloid population used in ATAC-seq and *in vitro* clonal differentiation in methylcellulose. E) UMAP of BM cells in the myeloid trajectory colored by metacluster (left) or projected FACS myeloid population (right). F) Histogram of normalized Tn5 insertion profiles centered at transcription start sites (TSSs) for the indicated ATAC-seq libraries. G) Barplots of normalized TSS insertion scores across biological and technical replicates. H) Histogram of insertion counts by fragment length for indicated ATAC libraries. I) UMAP of scATAC from healthy human BMMNCs colored by projections of biological and technical replicates of ATAC-seq libraries.

**Figure S5: Additional analysis of mPB and BM – related to Fig. 5**

A) Dotplot quantifying frequency of CD38- cells within CD34+ compartment by tissue/product. P-value derived from Wilcoxon rank sum test. B) Violin plots of marker expression by tissue/product. Diamond indicates median. C) Heatmap of median expression of metaclusters (top row colors), separated by tissue/product (second row colors).

**Figure S6: FL is comprised of cells with phenotypic profiles unique from other hematopoietic tissues and products – related to Fig. 6**

A) Barplot of frequency of CD34+ cells out of CD45+ cells in FL, separated by donor. B) UMAP of mononuclear CD45+ FL cells, colored by metacluster. C) UMAP colored by marker expression. D) Heatmap of median expression of leiden clusters, colored by metacluster annotation (top row). E) Frequency of FL metaclusters out of CD45+ cells. Dots indicate individual donors. Crossbar indicates mean.
